# Pharmacological proximities in the GPCR family discovered using contact-informed amino-acid and binding pocket similarities

**DOI:** 10.64898/2026.05.02.720972

**Authors:** Sean S. So, Tony Ngo, Andrey V. Ilatovskiy, Angela M. Finch, R. Peter Riek, Ruben Abagyan, Nicola J. Smith, Irina Kufareva

**Affiliations:** Orphan Receptor Laboratory, Department of Pharmacology, School of Biomedical Sciences, Faculty of Medicine & Health, UNSW Sydney, Sydney, NSW, Australia; Victor Chang Cardiac Research Institute, Darlinghurst, Sydney, Australia; Department of Biochemistry and Molecular Biology, Monash Biomedicine Discovery Institute, Monash University, Clayton, VIC, Australia; Skaggs School of Pharmacy and Pharmaceutical Sciences, University of California San Diego, La Jolla, CA, USA; Third Element Bio, San Diego, CA, USA; Health Sciences South Carolina, Columbia, SC, USA; School of Biomedical Sciences, Faculty of Medicine & Health, UNSW Sydney, Sydney, NSW, Australia

## Abstract

Understanding protein proximities in the theoretical ligand space is essential for developing therapeutics with desirable polypharmacology, predicting off-targets, and discovering surrogate ligands for poorly characterized proteins. This is especially important for G protein-coupled receptors (GPCRs) - a major class of drug targets, many of which still lack known ligands. Circumventing this limitation, we present GPCR-CoINPocket v2, a contact-informed metric for detecting GPCR pharmacological similarities from amino-acid sequences alone. We first establish a “gold standard” of pharmacological relatedness using ChEMBL-derived ligand sets. We then replace traditional evolutionary amino acid similarity matrices with a chemically-informed matrix derived from protein:ligand interaction patterns across 3,306 structures, significantly improving early detection of shared pharmacology between distantly homologous receptors. An additional unconstrained, contact-informed matrix further enhances predictive performance. Pilot application of the method revealed previously unrecognized similarities between the β_2_ adrenoceptor and three Class A peptide GPCRs, which we confirmed experimentally by demonstrating the binding of select ligands of these receptors to the β_2_. Dimensionality reduction of similarity scores recapitulates known receptor relationships and predicts neighbors of orphan GPCRs later confirmed experimentally. Overall, GPCR-CoINPocket v2 provides a powerful sequence-based framework to prioritize ligand space, predict polypharmacology, and accelerate GPCR drug discovery and deorphanization.

## Introduction

G protein-coupled receptors (GPCRs) have remained amongst the most highly clinically-targeted protein families to date^1^. Despite widespread interest as clinical targets, ~80 Class A (rhodopsin-like) GPCRs remain “orphans” unpaired to any cognate ligand, and often any ligand at all^2^. Given the pharmacological tractability of GPCRs, such receptors represent untapped sources of pharmacological intervention.

Unfortunately, the deorphanization rate of GPCRs in recent years is at an all-time low^3^. This does not reflect an absence of drug discovery efforts at these potential pharmacotherapeutic targets. Rather, this reflects limitations in current methods in breaking the orphan GPCR drug discovery loop: to identify a ligand, knowledge of signaling is typically required yet conversely, to interrogate signaling, a ligand is often needed^4^. The identification of a ligand can break this cycle, exemplified by the orphans GPR65/68^5^, MRGPRX2^6^, GPR139^7^, GPR182 (now deorphanized as ACKR5)^8–10^ and GPR25^11^.

Previously, we developed GPCR-Contact-Informed Neighboring Pocket (GPCR-CoINPocket), a metric of pharmacological similarity between Class A GPCRs intended to aid in these initial ligand selection steps^12,13^. The metric combined structural and phylogenetic data to encode ligand-binding information at the level of sequence similarity comparisons through weighting pairwise amino acid scores with ligand contact strengths observed in atomic-resolution Class A GPCR structures. For phylogenetically dissimilar (20-35% similarity) Class A GPCR pairs where sequence similarity is sufficient for high quality sequence alignment but phylogeny is no longer predictive of pharmacology, we observed that GPCR-CoINPocket indeed outperformed both transmembrane (TM) and binding pocket phylogeny in discriminating Class A GPCR pairs that share ligands from those that are pharmacologically unrelated.

Recent technological advances including AlphaFold 3^14,15^, Boltz-1^16^, and Boltz-2^17^ have streamlined the prediction of ligand-protein complex geometries and rapidly accelerated *in silico* drug screening. Yet, *in silico* discovery still faces key challenges: 1) it is unfeasible to predict the ligand geometries for even a miniscule fraction of the estimated 10^60^ members of the chemical universe^18^, even for a single receptor target, and 2) for validated hit compounds, it is nontrivial to predict their likely off-target binders. GPCR-CoINPocket addresses some of these challenges by providing pharmacological proximities between Class A GPCRs, both drastically minimizing the chemical search space and highlighting likely off-target receptors.

To improve the recognition of pharmacologically proximal receptor pairs with GPCR-CoINPocket, we turned to replacement of the underlying Gonnet mutational amino acid similarity matrix. Standard mutational matrices such as Gonnet’s matrix^19^, Dayhoff’s point accepted mutation (PAM) matrix^20^, and Henikoff and Henikoff’s block substitution matrix (BLOSUM)^21^ score sequence alignments by focusing on evolutionarily conserved features, including secondary structure conservation. While Gonnet’s matrix outperforms the other matrices in fold recognition^22^, structural information only vaguely encodes for chemical properties; thus, for predicting shared pharmacology, we instead propose the use of chemically- and/or contact-informed matrices that directly reflect ligand-binding properties to score pre-aligned sequences.

Here, we describe three distinct improvements to GPCR-CoINPocket that have hypothesized uses beyond application to GPCRs. We first developed a method to robustly classify proteins as pharmacologically related or unrelated based on chemical ligand sets derived from ChEMBL, and applied this to the classification of Class A GPCRs. We leveraged this classification to train an optimized amino acid similarity matrix informed by protein:ligand interaction patterns as derived from 3,306 liganded structures annotated in the Pocketome^23^ as a proxy for chemical information. When replacing the standard Gonnet matrix with our chemically-informed matrix, we reveal previously unappreciated relationships between the β_2_ adrenoceptor and various peptide receptors, and validate these findings with radioligand binding assays. Seeking to further improve the accuracy of GPCR-CoINPocket, we additionally trained an unconstrained contact-informed amino acid similarity matrix with near-perfect ability to discriminate pharmacologically similar from dissimilar Class A GPCR pairs. We summarized each-to-each comparisons with Uniform Manifold Approximation and Projection^24,25^ (UMAP), revealing pharmacological neighbors of orphan GPCRs that were only confirmed in recent years despite training on structural data from a cutoff of mid-2018. We hope that these advances will help minimize the chemical search space in ligand identification campaigns for orphan receptors, characterize polypharmacology of known and yet-to-be discovered drugs, and reduce the probability of unwanted side effects by identifying and prioritizing the likely secondary targets. We also believe that our matrices will illustrate the importance of considering how amino acid similarities are calculated, and aid in refining and training deep and traditional machine learning predictors of pharmacological similarity.

## Results

### A formal classification scheme defining pharmacologically similar or dissimilar proteins

To quantitatively assess the performance of GPCR-CoINPocket in identifying pharmacologically proximal receptor pairs, and to evaluate the impact of method modifications^13^ we first sought to design a generalizable yet empirical metric describing the pharmacological relatedness of Class A GPCRs. From ChEMBL v27, we extracted 85,734 GPCR:ligand pairs (encompassing 174 Class A receptors) with high quality binding data and submicromolar affinity (**Extended Data Fig. 1A** and **B**). To restrict analysis to only unique chemotypes shared between GPCRs and account for uneven distribution of ligand set sizes for each GPCR (**Fig. 1A, Supplementary Fig. 1**), ligand sets were hierarchically clustered by Tanimoto distance with a cutoff of 0.1. These unique chemotypes were used to construct asymmetric and pairwise measurements of the pharmacological relatedness between given pairs of GPCRs *i* and *j* such that each score, *S*_*ij*_ or *S*_*ji*_, represented the estimation of the number of ligands from receptor *i* acting at receptor *j*, and vice versa (**Fig. 1B**). We observed that *S*_*ij*_ and *S*_*ji*_ scores were heavily skewed towards fewer shared chemotypes (**Fig. 1B**) – an expected result given that most GPCRs are understudied and have few ligands in ChEMBL^4^.

**Fig. 1.**
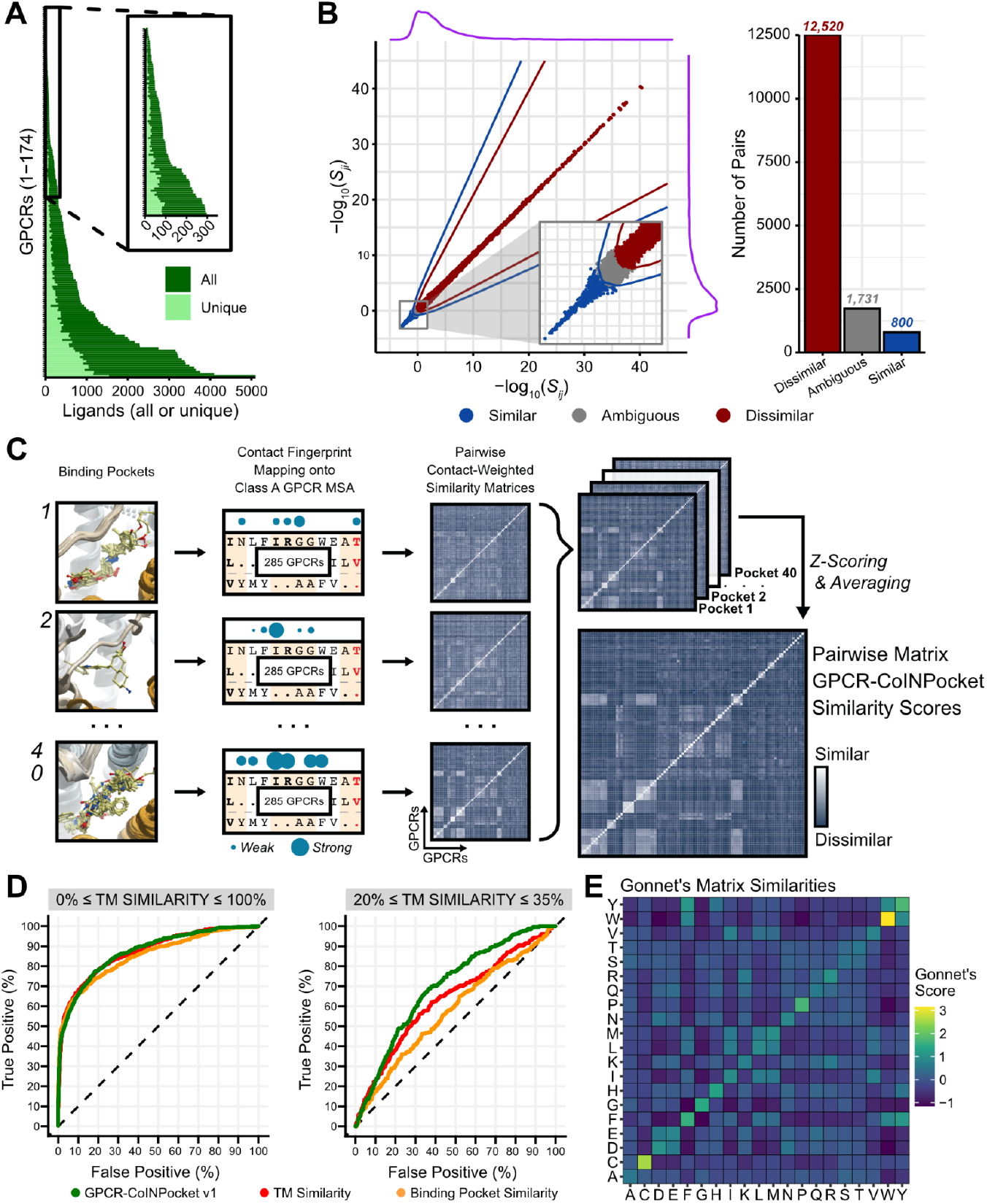
Empirical classification of pharmacologically similar and dissimilar GPCR pairs. **a**, Number of ligands or unique chemotypes for each Class A GPCR in ChEMBL v27. Inset, zoom of GPCRs with few ligands. **b**, Asymmetrical ligand scaffold set similarity measurements are used to define pharmacologically similar, dissimilar, or ambiguous pairs. Inset, zoom. Purple, density curves. **c**, Schematic of the CoINPocket methodology^13^ applied to Class A GPCRs. **d**, ROC curves comparing GPCR-CoINPocket v1 using updated ligand fingerprints, transmembrane (TM) similarity, and binding pocket similarity in distinguishing similar GPCR pairs from dissimilar ones across either all GPCRs or those with homology too low for accurate modeling but sufficient similarity for high accuracy sequence alignment. **e**, Heatmap representing pairwise amino acid similarity scores as defined by the Gonnet matrix^19^. MSA = multiple sequence alignment.

Conceptually, irrespective of ligand set sizes, if two proteins share a sufficient number of ligands between them, they should be considered pharmacologically related to some degree. Here, we defined pharmacologically similar receptors using a hyperbolic scoring function (see **Methods** and **Supplementary Note 1**) such that GPCR pairs with *S*_*ij*_ and *S*_*ji*_ scores approximately 5 or higher (interpreted as sharing approximately 5 unique chemotypes or more) were considered pharmacologically similar (**Fig. 1B**). Similarly, GPCR pairs with *S*_*ij*_ and *S*_*ji*_ scores approximating 1 or less (interpreted as sharing less than 1 unique chemotype) were considered pharmacologically dissimilar. Using the GPCR:ligand pairs extracted from ChEMBL v27, this resulted in 800 (5.32%) similar pairs, 12,520 (83.18%) dissimilar pairs, and 1,731 (11.5%) ambiguous pairs. These ambiguous pairs represented GPCR pairs with more shared compounds than would be classifiable as completely pharmacologically dissimilar, but still not enough empirical evidence to assume shared pharmacology. We assessed this classification metric by examining the pairwise Tanimoto distance distributions between ligand sets of GPCRs with high and low homology that were categorized as pharmacologically similar, dissimilar, or ambiguous (**Extended Data Fig. 2**), and found the scheme to reflect literature regarding these receptor pairs^26–29^.

We next tested whether the metric could be used to assess the performance of pharmacological relatedness predictors. We benchmarked the ability of GPCR-CoINPocket (described in **Fig. 1C**) transmembrane similarity, and binding pocket similarity to discriminate pharmacologically related receptors from unrelated ones using a receiver-operator characteristic (ROC) curve (**Fig. 1D**). These metrics were equally competent at identifying pharmacologically similar pairs across all GPCRs, consistent with our expectations that both TM and binding pocket sequence similarity are predictive of pharmacological similarity for highly homologous receptors. For distantly related GPCRs (20% to 35% homologous) where homology is not predictive of pharmacological similarity, GPCR-CoINPocket outperformed both TM and binding pocket sequence similarity. These results closely match our previously published work^13^ indicating that the classification scheme presented here is robust to changes in ChEMBL versions, despite large shifts in GPCR:ligand pair confidence scores, assay type assignation, and unique GPCR:ligand pairs between updates (**Extended Data Fig. 1**).

### Improvements to the GPCR-CoINPocket metric

Having formalized an intuitive understanding of what constitutes pharmacological similarity into an empirically-derived scheme for pharmacological relatedness, we next sought to improve the performance of GPCR-CoINPocket in identifying receptor pairs with shared pharmacology. In brief, the CoINPocket approach projects structurally observed ligand interaction fingerprints from the Pocketome^23^ from a given pocket across all Class A GPCRs in a multiple sequence alignment, and weights each positional pairwise sequence similarity by the distance approximation of observed GPCR:ligand contact strengths^13^ For each structurally-observed orthosteric pocket (defined by the GPCR:ligand interactions), these scores are summed for each GPCR pair, then z-scored across the set. Summed and normalized similarities are averaged across all pockets to form a GPCR-CoINPocket score for each GPCR pair (**Fig. 1C**).

Here, we sought to improve the GPCR-CoINPocket metric by optimizing both the structural and sequence components of the algorithm. We did not observe an improvement in the predictive power of GPCR-CoINPocket v1 despite a large increase in structural data between our previous work and Pocketome v18.04^23^ (**Extended Data Fig. 3**), likely because the new ligand binding fingerprints largely matched those previously incorporated (**Extended Data Fig. 4**). Instead, we focused on replacing Gonnet’s mutational matrix (**Fig. 1E**), designing a metric that reflected amino acid similarity in terms of functional group-binding propensities instead of mutational rates and evolutionary fold conservation.

### Development and application of an amino acid similarity matrix reflecting structurally-derived ligand recognition patterns

Comparisons of how frequently or strongly (and thus, preferentially) individual amino acids interact with certain chemicals or ligands over others is empirically reflective of ligand recognition patterns. To describe amino acids in these terms, we obtained vectors of amino acid:ligand binding frequencies from all 3,306 liganded proteins in Pocketome v18.04. To ensure that interaction vectors were comparable across residues and pockets, ligands were fragmented into ChEMBL-mined bioisostere groups common across the chemical universe (**Fig. 2A**). Interaction strengths between every amino acid and each of the mined 83,043 chemical fragments were measured in all 3,306 protein:ligand structures. Interaction strengths were approximated by a weighted inverse distance metric^30,31^ and classified as polar, non-polar, or backbone depending on the interacting atoms of the amino acid using BaSiLiCo ^32^ and as previously described^23,30,33^. Contact interaction strengths were recorded on a positional basis whilst non-interaction frequencies were recorded on a per-pocket basis in an attempt to reduce skew (for every contact a fragment makes with an amino acid, it does not make it with up to 19 others). To ensure contact vectors were sufficiently sized for pairwise amino acid comparisons, interaction vectors were considered only if more than 10 contacts across the Pocketome were observed. The resultant 3,984 fragments were further aggregated by hierarchical clustering of Tanimoto distances, generating 1,233 fragment clusters (**Fig. 2A**). Thus, each amino acid is described by interaction vectors (including non-contact frequency and contact strengths) from each of the 1,233 fragment clusters for non-polar, polar, and backbone interaction types, provided as **Supplementary Data 3**.

**Fig. 2.**
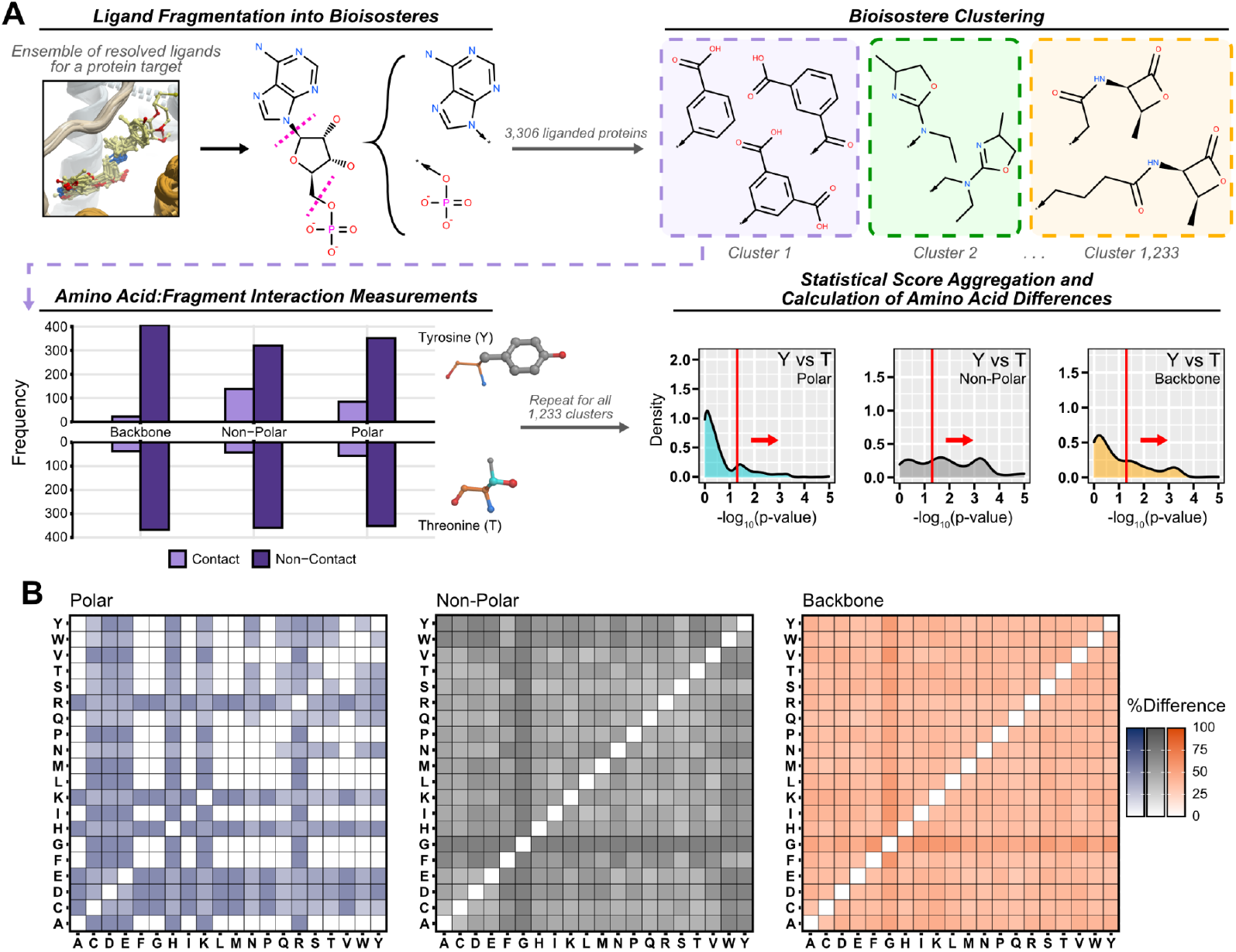
Generation of residue:ligand chemically-informed amino acid matrices for each contact type. **a**, Schematic of the data collection strategy for describing amino acid similarities in chemical terms. Liganded structures from the Pocketome v18.04^23^ were substructure-matched into bioisostere fragments (conceptual “cut sites” shown as pink dashed lines), then filtered and clustered by Tanimoto distance. Contacts were measured and non-contacts quantified across each of the 3,306 liganded pockets. Aggregated contact vectors were used for statistical measurement. The distribution of 1,233 p-values (one for each cluster) was plotted for each pairwise amino acid comparison, for each contact type, and the AUC of the kernel-smoothed densities of p-values ≤ 0.05 estimated the amino acid differences. An example of Y (Tyr, tyrosine) vs. T (Thr, threonine) is shown, along with a colored 3D representation of their chemical structures: gray, non-polar regions; orange, backbone regions; cyan, polar regions, red, acidic groups; blue, basic groups. **b**, The amino acid contact-informed difference matrices for each contact type.

We then assessed whether amino acids differed in their propensity to bind particular chemical clusters or whether one interacted more strongly than another using chi-squared and non-parametric Mann-Whitney/Wilcoxon rank-sum tests. These tests were performed for each of the 1,233 clusters against every amino acid pair, for each contact type (**Fig. 2A**). The AUC of the kernel-smoothed distributions of the log_10_-scaled significant p-values (p≤0.05) defined the degree of difference between each amino acid pair (**Supplementary Data 1**), and are represented for each contact type as heatmaps of pairwise, contact-type similarities/differences (**Fig. 2B**). Many of the pairwise amino acid scores are consistent with intuitive expectations for their ligand binding preferences; for example, aromatic residues are most similar to each other and are dissimilar to other non-polar residues in the non-polar matrix (**Fig. 2B**).

Application of these chemically-informed pairwise amino acid matrices to GPCR-CoINPocket required the assembly of the three contact types into a single value for each amino acid pair. We imagined a simple linear combination of the three contact types on a fractional 0 to 1 scale, where each contact type was weighted by a coefficient, and all coefficients were positive and summed to 1 (**Fig. 3A**). To determine the optimal combination of these parameters, the gradient-free search methods COBYLA (constrained optimization by linear approximation) and Nelder-Mead simplex were employed. The algorithms sought to maximize the AUC of a square-root-scaled ROC curve that prioritized early detection of pharmacologically similar pairs in the low homology (20% to 35%) GPCR pair subset (**Fig. 1D**). To identify the global maxima, non-polar and polar weights were initialized in a grid-start manner from all combinations of 0 to 1 in steps of 0.1.

**Fig. 3.**
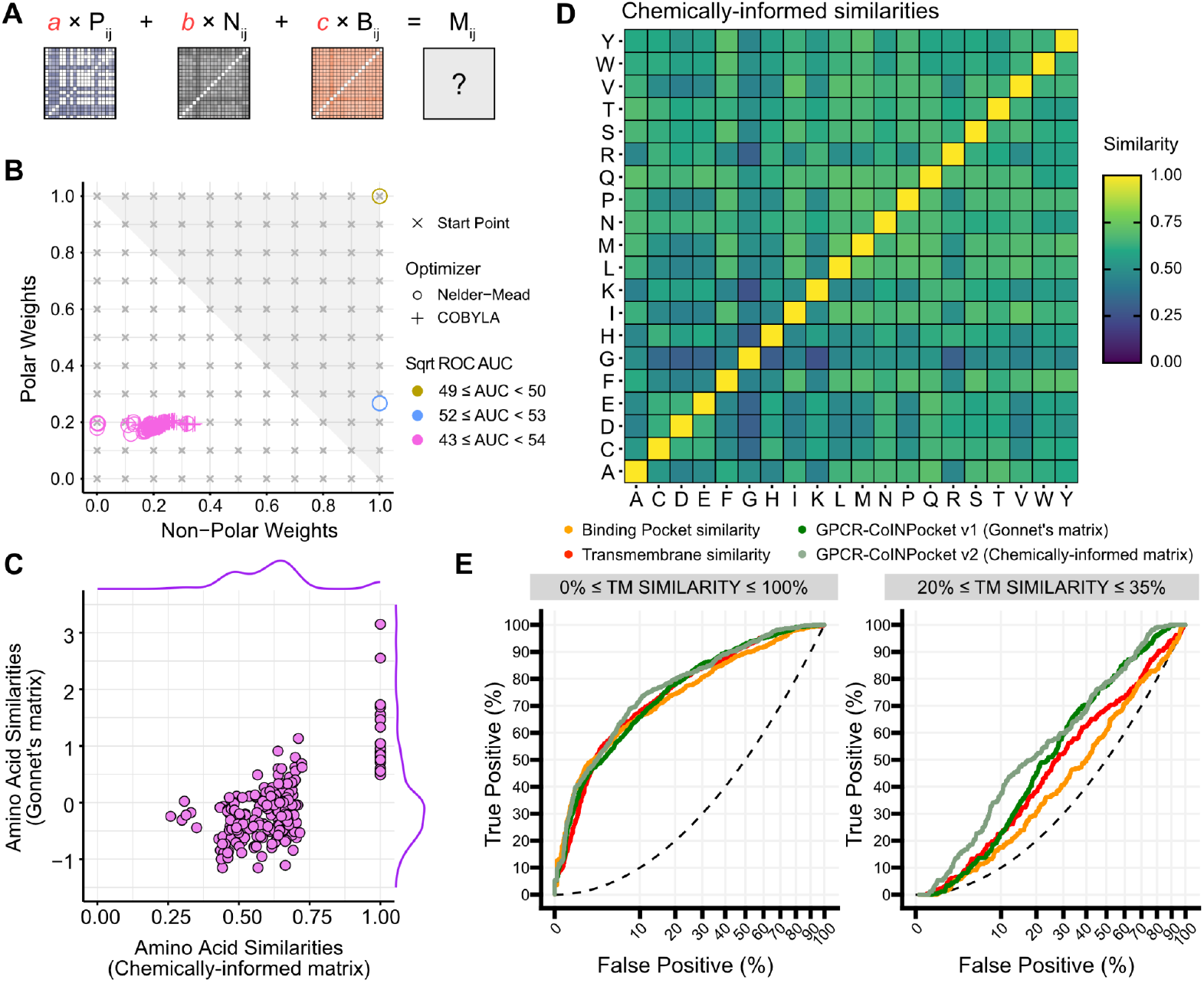
Optimization and application of a chemical-contact-informed amino acid similarity matrix. **a**, A schematic showing the coefficients (red) that are optimized by the COBYLA or Nelder-Mead simplex algorithms to maximize early detection AUC. **b**, A scatterplot of the final minimized weight coefficients (non-polar, polar) exposed to the optimizers. Gray crosses represent all start points. Pluses and open circles represent optimized weights for COBYLA and Nelder-Mead simplex, respectively. The color of symbols represents the early-detection AUC of the final weight triplet including derived backbone weights (1 - sum of polar and non-polar): gold, 49 to 50; blue, 52 to 53; pink, 53 to 54. The gray triangle represents disallowed regions where backbone weights are negative. **c**, A scatterplot comparing amino acid pairwise similarity scores for the chemically-informed matrix or Gonnet’s mutational matrix. Each point represents a single amino acid pair. Marginal density plots (top, right) show the distribution of data along each axis. **d**, Pairwise amino acid similarity scores for the chemically-informed matrix are shown as a heatmap. **e**, Comparison of GPCR-CoINPocket v1, GPCR-CoINPocket v2 with chemically-informed matrix, transmembrane (TM) similarity, and binding pocket similarity in distinguishing true pharmacological pairs from dissimilar pairs across all GPCRs or those with low but sufficient similarity for high accuracy homology modeling. The x-axis is represented on a square-root scale to emphasize early detection events. Pharmacological similarity definitions are as derived from ChEMBL v27.

Within these constraints, both COBYLA and Nelder-Mead converged around three principal points with correspondingly different early-recognition-weighted ROC curve AUCs (**Fig. 3B**). Of these, only the best-performing weight coefficients (highest AUC) satisfied all constraints. The median value of the coefficients with scaled ROC curve AUCs between 53 and 54 was 0.19, 0.2, and 0.61 for polar, non-polar, and backbone matrices respectively. Using these coefficients, we defined a final amino acid similarity matrix, empirically derived from observed residue:ligand contacts (**Fig. 3D**). This chemically-informed matrix naturally sets self-to-self comparisons to 1 unlike the Gonnet matrix, which considers mutational and evolutionary frequencies (**Fig. 3C**), and the pairwise amino acid scores are provided in **Supplementary Data 1**.

GPCR-CoINPocket using the chemically-informed matrix (**Fig. 3D**) outperforms the implementation with Gonnet’s matrix by approximately 2-fold in early detection up to 15% false positives with no loss to overall predictive power (AUCs 71 vs 72, respectively; **Fig. 3E**). Of the top 10% highest scoring distantly related GPCR pairs (679 out of 6784), 91 are pharmacologically similar (as defined above and in **Fig. 1**), while only 56 pairs were captured by the implementation using Gonnet’s matrix. Despite only training on low homology pairs, we also observed improved predictive performance across all GPCRs (**Fig. 3E**). Further, optimization using a 5-fold stratified cross-validation strategy resulted in similar optimized weights, suggesting that these weights were stable (**Extended Data Fig. 5**). Overall, this suggests that the matrix was not heavily overtrained to recognize only early-ranking distantly related Class A GPCR pairs. We term the GPCR-CoINPocket implementing the chemically-informed amino acid similarity matrix developed here GPCR-CoINPocket v2 and supply all calculated pairwise GPCR comparisons as **Supplementary Data 2**.

### Identification of secondary activities of peptide receptor-targeting small molecules at the β_2_ adrenoceptor

To prospectively validate GPCR-CoINPocket v2, we asked whether it could identify pharmacological neighbors even for very well-characterized receptors, leading to the discovery of previously unknown off-target effects for ligands of these receptors. When ranking Class A GPCRs by their TM similarity to the β_2_ adrenoceptor, aminergic GPCRs ranked at the top of the list (**Fig. 4A**), consistent with the high level of ligand^34^ and sequence conservation within the family^35–37^. However, when ranking receptors by GPCR-CoINPocket v2 score, 15 peptide GPCRs ranked above even some aminergic receptors (**Fig. 4A**). These included members of the somatostatin, melanin-concentrating hormone, vasopressin, oxytocin, opioid, and urotensin receptor families (**Supplementary Data 4**).

**Fig. 4.**
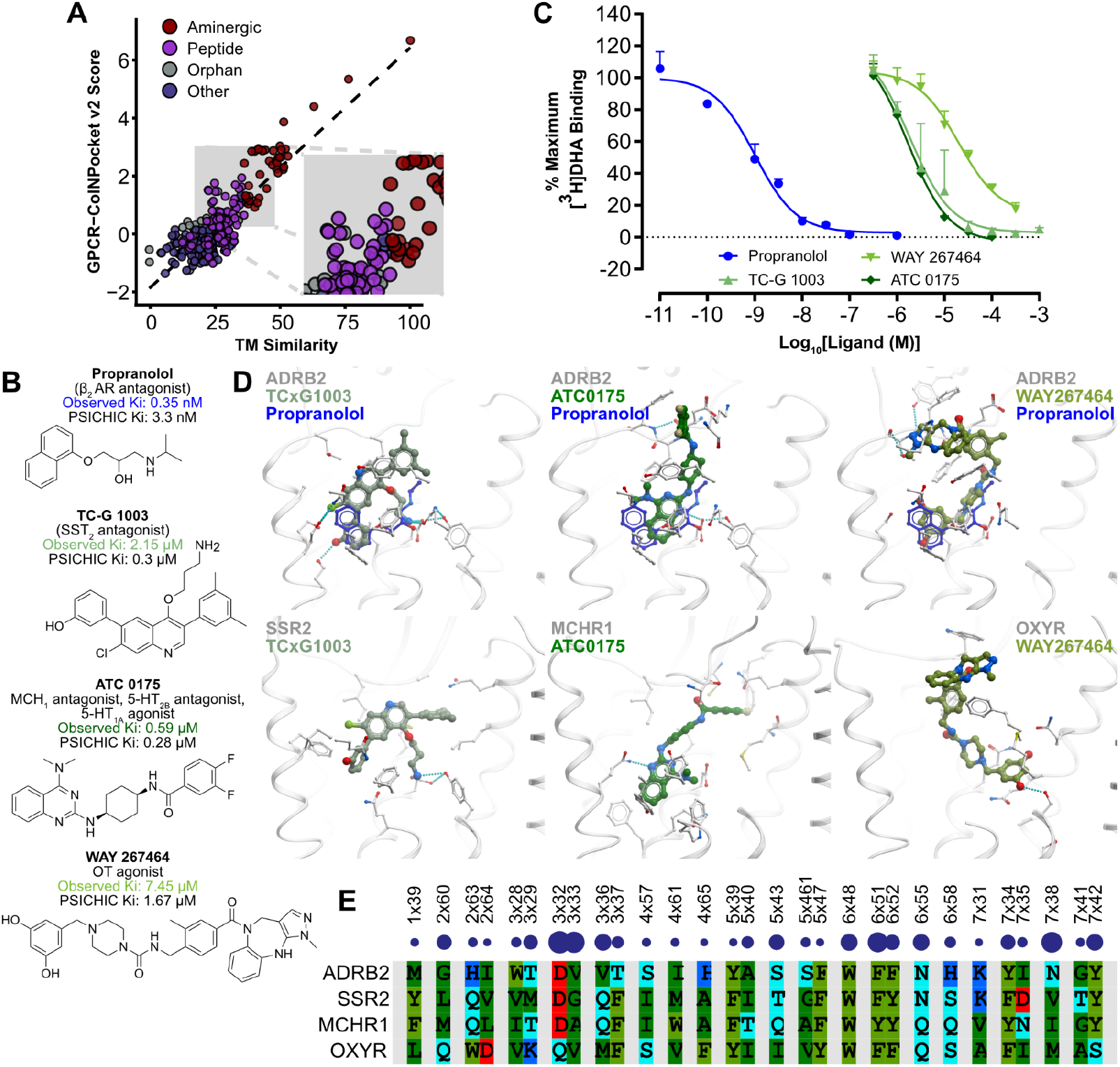
GPCR-CoINPocket v2 identifies unexpected neighbors of the β_2_ adrenoceptor. **a**, The pharmacological neighborhood of the β_2_ adrenoceptor. The transmembrane sequence similarity of 285 Class A GPCRs to the β_2_ adrenoceptor are plotted against their corresponding GPCR-CoINPocket v2 scores. GPCRs are coloured broadly by classification: aminergics (red), peptide (purple), orphans (gray), and others (navy). **b**, Chemical structures of micromolar affinity ligands assessed in this study. Observed K_i_ (affinity) of experimental ligands (green) or propranolol (blue) from competitive radioligand binding are displayed above each structure along with the PSICHIC-predicted K_i_. **c**, Competitive radioligand binding assays using putative surrogate ligands. Binding assays used crude purified membranes of COS-1 cells transiently transfected with β_2_ adrenoceptor. Unlabelled competitor ligands were titrated against 0.5 nM [^3^H]DHA and their affinity derived. Points and bars represent means ± SEM of n=3 experiments performed in triplicate. Curves were fit to a logistic single site binding model using GraphPad Prism v9. **d**, Boltz-1 and AlphaFold 3-predicted binding poses of propranolol and the low micromolar affinity off-target ligands co-folded with the β_2_ adrenoceptor and their cognate receptors. In blue, the binding pose of the β_2_ antagonist propranolol is shown bound to the β_2_ adrenoceptor.

To validate these predictions, we assessed the competitive binding affinity between the β_2_ adrenoceptor and 8 diverse small molecule ligands of 6 peptide GPCRs (somatostatin receptor SST_2_, somatostatin receptor SST_4_, neuropeptide S receptor NPS, melanin-concentrating hormone receptor MCH_1_, orexin receptor OX_2_, and oxytocin receptor OT) (**Fig. 4B**). In competitive radioligand binding assays, modulators of the OT (WAY 267464), 5-HT_1A_/5-HT_2B_/MCH_1_ (ATC 0175), and SST_2_ receptors (TC-G1003) had low micromolar affinity for the β_2_ adrenoceptor (**Fig. 4C**); all others tested had unresolved binding curves or were inactive at micromolar concentration (**Supplementary Fig. 2, Supplementary Data 4**). For ATC 0175 and WAY 267464, radioligand dissociation kinetic assays further supported the likelihood that the binding was orthosteric (**Supplementary Fig. 2**). To better understand how these compounds might bind to the β_2_ adrenoceptor, we applied Boltz-1^16,38^ and Alphafold 3^14^ to predict ligand binding poses by undirected co-folding of the β_2_ adrenoceptor and the hit compounds (**Fig. 4D**). The orthosteric pocket was the most energetically favorable binding location for these predicted receptor:ligand complexes. When aligning the β_2_, MCH_1_, OT, and SST_2_ receptors by the important Class A GPCR ligand contacts, defined by the consensus interaction fingerprint across all Class A binding pockets (**Extended Data Fig. 4**) we found a high degree of sequence similarity between these receptors (**Fig. 4E**). Moreover, the compounds appeared to engage via the most prototypical interactions in the β_2_ adrenoceptor (**Extended Data Fig. 6**), further supporting orthosteric binding of the hit compounds at this receptor. Overall, we identified 3 peptide receptor ligands with moderate-to-low affinity at the β_2_ adrenoceptor, highlighting previously unappreciated pharmacological similarities between the cognate receptors of these compounds and the β_2_ adrenoceptor, as predicted by GPCR-CoINPocket v2.

To highlight the utility of GPCR-CoINPocket in minimizing the chemical search space for *in silico* screening, we submitted our proposed ligand:GPCR pairings to the PSICHIC server^39^, a state-of-the-art graph-based affinity prediction model that generates a predicted binding affinity and functional mode (agonist, antagonist, non-binder). We found the predictions to be relatively accurate for our findings (**Supplementary Data 4**): of the 3 compounds we found to bind to the β_2_ adrenoceptor, PSICHIC was able to accurately predict affinity (within 0.5 log units) and the functional mode, though it was ambiguous for its prediction of WAY 267464 (35% non-binder). Amongst the remaining 5 compounds we examined, PSICHIC also accurately predicted non-binding for L-368,899 (82% non-binder) and agreed with our initial prediction of possible binding for the other 4 compounds (0-1% non-binder), for which it predicted binding affinities similar to its prediction of the β_2_ antagonist propranolol. Overall, PSICHIC displayed great utility in accurately predicting compound activity and binding likelihood. We propose a framework for future *in silico* studies whereby GPCR-CoINPocket is used to minimize the chemical search space for ligand picking, compounds are ranked by their predicted binding affinity, and predicted compounds are tested *in vitro* to confirm predictions.

### Development and application of an unconstrained contact-informed amino acid similarity matrix

Having established, both retrospectively and prospectively, the validity of GPCR-CoINPocket v2 with a chemically-informed matrix, we next asked if we could further improve the recognition of receptor pharmacological similarities and neighborhoods by relaxing the amino acid comparison matrix optimization constraints and allowing individual scores to vary independently. To this end, we used the Nelder-Mead simplex method to directly optimize matrix elements with the goal of maximizing the same ROC AUC as before, but naive to the chemical matrices previously described (**Fig. 2B**). Scores were allowed to vary between −1 and 1, with the intention that dissimilarity (−1) should impose a penalty on the score, and that this differed from indifference (0).

As with the chemically-informed matrix, a range of AUCs was observed, though AUCs tended to be much higher, between 65 and 82 (**Extended Data Fig. 7**). Unexpectedly, the top performing optimization runs with square-root-weighted ROC curve AUCs above 80 all arose from initializations where all amino acid pairs but one were initialized at ~1.0 (**Extended Data Fig. 7**). From these optimization runs, the median optimized value for each amino acid pair was taken to be the final optimized weight (**Fig. 5A**). Converged amino acid similarities for the unconstrained contact-informed matrix varied between 0.1 and 1.0, and were not correlated with the chemically-informed or Gonnet’s matrices (**Fig. 1E, 3D** and **5B**). The optimized amino acid scores broadly mirrored our expectations regarding chemical contacts: for example, non-polar amino acids were most closely related to each other, as were polar amino acids. However, we noted a surprising lack of similarity between arginine and histidine, reflecting their distinct roles in recognition of chemical compounds, and contrasting their high similarity in the context of protein folding. When applied to GPCR-CoINPocket, the unconstrained contact-informed matrix vastly outperformed all other metrics in the recognition of pharmacologically related but phylogenetically distant GPCR pairs according to ChEMBL v27 definitions (**Fig. 5C**). Similar improvements were observed when considering all GPCR pairs as a whole and predictive accuracy was consistent across ChEMBL versions 21, 27, and 35 (**Extended Data Fig. 1C**) despite sharing just 52.7% of unique ligand:receptor pairs between them (**Extended Data Fig. 1B**), suggesting the model was not overtrained to recognize only phylogenetically distant pairs present in ChEMBL v27.

**Fig. 5.**
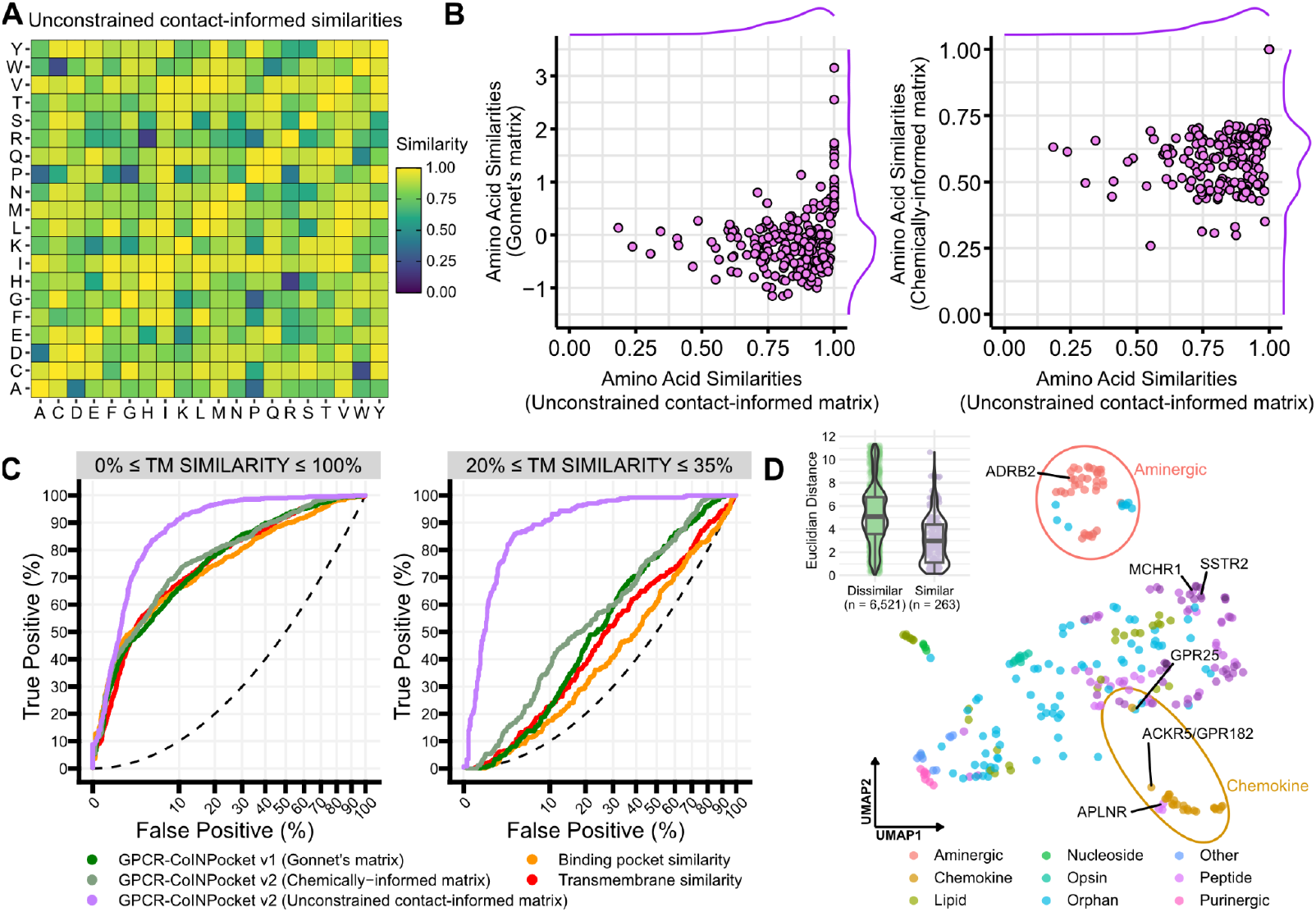
Optimization and retrospective application of an unconstrained contact-informed amino acid similarity matrix. **a**, Pairwise amino acid similarity scores for the unconstrained contact-informed matrix are shown as a heatmap. **b**, A scatterplot comparing amino acid pairwise similarity scores between the unconstrained contact-informed matrix to the chemically-informed or Gonnet’s mutational matrices. Each point represents a single amino acid pair. Marginal density plots (top, right) show the distribution of data along each axis. **c**, Comparison of GPCR-CoINPocket v1 and v2 with either the chemically-informed or unconstrained matrices, transmembrane (TM) similarity, and binding pocket similarity in distinguishing true pharmacological pairs from dissimilar pairs across all GPCRs or those with low but sufficient similarity for high accuracy sequence alignment. **d**, UMAP representation of the global Class A GPCR family considered in this study. Each point represents a single Class A GPCR, with similarity vectors provided in **Supplementary Data 2**. Families are colored as shown. Receptors of interest have been labelled, and the chemokine and aminergic families have been circled. The inset shows the distribution of all distances between every point, grouped by pharmacologically similar (lavender) or dissimilar (green). GPCRs are named in accordance with UniProt-SwissProt protein names.

To examine the ability of GPCR-CoINPocket with the unconstrained contact-informed matrix to infer pharmacological similarities on a family level, we represented Class A GPCRs with Uniform Manifold Approximation and Projection (UMAP) clustering^24,25^. Receptor subfamilies generally clustered in ways that recapitulated our understanding of Class A GPCR pharmacological landscape (**Fig. 5D**). To estimate the performance of UMAP embedding, pairwise GPCR distances in the UMAP space were binned according to pharmacological similarity ground truths derived from ChEMBL ligand sets. We observed that UMAP embedding more frequently placed similar receptors together and dissimilar receptors further apart, reflecting empirical ligand-centric relationships, despite being derived purely from sequence.

Closer inspection yielded important differences compared to representations of Class A GPCRs binding-pocket phylogeny dendrograms^13,34,37^. For example, GPR182 is phylogenetically positioned closest to the apelin, GPR25, and GPR15 receptors, and with greater similarity to the formyl peptide receptors than to CC or CXC chemokine receptors^13^. However, in November 2024, GPR182 was deorphanized as an atypical receptor for chemokines^8,9^ (and renamed as ACKR5). Our CoINPocket v2-based UMAP representation places ACKR5 within the chemokine receptor cluster and proximal to apelin and related receptors with which it also shares ligands^8^, despite training on data available pre-2018. GPR25, another orphan GPCR, was also placed next to XCR1, consistent with its recent deorphanization as a receptor for CXCL17^11^, another ligand for ACKR5^10^. Furthermore, we observed the peptide GPCRs to broadly group together, with the somatostatin and melanin-concentrating hormone receptor subfamilies located closest to the biogenic amine subfamily, supporting our earlier findings (**Fig. 4**). The complete set of predicted pairwise GPCR similarity scores using various metrics, with pharmacological similarities and dissimilarities as defined by ChEMBL versions v21, v27, and v35 is provided as **Supplementary Data 2**. Experimental validation of these newly predicted neighborhood relationships and further optimization of the amino acid matrix including application to deep learning approaches is beyond the scope of this work and will be the subject of future studies.

## Discussion

Here, we present GPCR-CoINPocket v2, an update to our previous metric of Class A GPCR pharmacological similarity^13^. Substantial improvement in early predictive capacity was attained by replacement of the Gonnet amino-acid similarity matrix with a contact-informed matrix derived from protein:ligand complexes. Even greater retrospective improvement was achieved with the use of a purely contact-informed and unconstrained contact-informed matrix, though these new pharmacological neighborhoods remain to be experimentally validated. Focusing on the β_2_ adrenoceptor we identified ligands of the SST_2_, MCH_1_, and OT receptors that displayed low micromolar off-target affinity to the β_2_ adrenoceptor. In doing so, we demonstrate that GPCR-CoINPocket is able to recapitulate ligand features at the level of amino-acid sequence and predict nontrivial shared ligand activities between phylogenetically distant receptors.

Predicting receptor pharmacology from sequence alone is inherently difficult. Early methods have demonstrated some level of success at encoding ligand information at the level of sequence by examining the most highly conserved residues within a GPCR subfamily, with a rationale that critical ligand-binding amino acids would be less subject to mutation^37^. Yet, attempts to generalize this across GPCR subfamilies by assessment of phylogeny at the level of binding pocket sequence similarity still failed to capture known relationships between GPCR targets^34^. With GPCR-CoINPocket, we have captured near-ligand-centric resolution of pharmacological neighborhoods using sequence alone, by supplementation of sequence similarities with conserved Class A GPCR binding patterns and amino acid interaction propensities.

Given the resolution of the CoINPocket approach, its applications are broad. By identifying pharmacological neighbors target proteins, it is possible to predict where ligands may display off-target activity. This can provide insight into unexplained clinical off-target effects, or be useful in pre-clinical pipelines for rational early counter-screening of lead compounds against undesirable but CoINPocket-predictable off-target vulnerabilities. Conversely, it may also be used to identify protein groups with *desirable* shared polypharmacology for multi-targeted approaches. Importantly for orphan GPCR drug discovery and *in silico* screening, CoINPocket may also aid in identifying liganded neighbors of orphans as a source for surrogate ligands^4,11^, as we have retrospectively seen for GPR25^11^ and ACKR5^8–10^. This enables focusing of the search space for both chemical discovery and deorphanization. Notably, application of the CoINPocket approach towards a non-GPCR family, lysine methyltransferases^40^, has also yielded first-in-class inhibitors of the orphan oncoprotein, NSD1, a highly relevant cancer drug target^41,42^.

There are limitations with the current iteration of GPCR-CoINPocket (and the broader CoINPocket approach) worth noting. Despite the demonstrated success of the CoINPocket approach in discriminating pharmacologically related proteins from unrelated ones^4,11^, the contacts that function in conjunction with the matrices require alignable pockets (and therefore knowledge of binding sites). Broadly, this means that predictions of off-target activity are limited to within the alignable protein family of interest. This is in direct contrast with ligand-focused metrics of pharmacological similarity^34,43,44^ that are capable of identifying off-target neighbors from entirely unrelated proteins, as has been shown for the amebicide emetine, which exerts its action partially via 40S ribosomal subunit binding^45^ and displays micromolar off-target antagonism of α_2_ adrenoceptors^43^. For GPCR-CoINPocket, this limitation is somewhat an advantage: low or absent ligand diversity is no longer an obstacle in measuring pharmacological proximities for orphan receptors. However, to mitigate the limitation of only detecting off-target activity between protein family members, we envision that further applications of the CoINPocket approach will move beyond protein families and focus on structurally alignable domains or motifs with ligand-binding functions. This will be the subject of future research.

Modern structure-based drug discovery campaigns for understudied GPCRs^46^ now commonly leverage deep-learning-assisted approaches such as ligand:protein complex co-folding^14–17,38^ and affinity prediction^39^, with geometrical accuracies approaching or even exceeding that of low-resolution experimental complexes ^46,47^. Yet, *in silico* screening campaigns against challenging targets such as orphan GPCRs still face substantial challenges. Most crucially, for a target with unknown pharmacology, it is not simple to understand the chemical space in which its ligands reside; even the readily-synthesized and accessible chemical space (e.g. Enamine REAL Space), contains over 94 billion unique molecules. Thus, for successful *in silico* screening campaigns, methods are needed to minimize the search space and simultaneously provide likely off-target neighbors for cross-validation of target selectivity. While deep-learning-accelerated pipelines now exist to attempt to broadly sample the chemical universe^48^, we hypothesize that success might be more easily achieved by deeply sampling pharmacological spaces where ligands are likely to exist; the so-called “low-hanging fruits”. Here, the CoINPocket approach has demonstrated an ability to resolve previously unknown pharmacological neighborhoods, though prospective direct application to orphans remains the goal of future research.

Ultimately, by translating structural interaction patterns into amino acid similarity matrices reflective of real-world ligand binding patterns, GPCR-CoINPocket provides a robust sequence-based framework to navigate intractable chemical landscapes, predict pharmacology, and streamline *in silico* drug discovery.

## Online Methods

### Data Acquisition and Chemoinformatics

#### Ligand fingerprinting and chemical distances

In this study, all Tanimoto distances between ligands were calculated using Molsoft Internal Coordinate Mechanics (ICM) v3.9.1a using proprietary fingerprint definitions. They include feature definitions for linear chains up to 7 atoms in length, non-linear components, and ring components.

#### Extraction of GPCR ligand sets from ChEMBL

Ligand structures and associated binding affinity data for all Class A GPCRs for ChEMBL versions 21, 27, and 35 were extracted from locally-hosted MySQL ChEMBL databases (https://chembl.gitbook.io/chembl-interface-documentation/downloads). To prevent contamination with potential off-target binders, compounds were only considered if they had pChEMBL activity values (a surrogate measure for K_D_ or K_i_) ≥ 6 in binding assays with confidence scores of 9 as rated by ChEMBL curators (the highest available), without known validity issues. Where duplicate values were observed, the 80th percentile value was taken as a conservative approximate of ligand activity. To maximize confidence in ligand set comparisons, ligand sets for receptors with less than 10 ligands in the data set were not considered.

#### Cross-version ChEMBL summary comparisons

To summarize major differences between ChEMBL versions, the full list of ligand sets including their cognate receptors were compared. Specifically, the overlap between the number of all ligand:receptor pairs, the number of unique receptors, and the number of unique chemotype:receptor pairs was assessed. To generate the Venn diagrams in **Supplementary Fig. 1A**, *ggvenn()* as implemented in the *ggvenn v0*.*1*.*10* R library was used.

### Pharmacological Similarity Scoring and Benchmarking

#### Calculation of pharmacological similarity scores between Class A GPCRs

The calculation of pharmacological similarity between a given pair of GPCRs *i* and *j*, with their associated ligand sets *L*_*i*_ and *L*_*j*_, was as follows. Ligand sets were first clustered by bottom-up hierarchical clustering using the Unweighted Pair Group Method with Arithmetic Mean (UPGMA) algorithm as implemented in ICM, with clusters defined by a Tanimoto distance cut-off of 0.1. The Tanimoto distances between every ligand cluster from *L*_*i*_ and *L*_*j*_ was calculated, and vice versa. With the understanding that each cluster represented a unique chemotype, the number of chemotypes from *L*_*i*_ for which the closest chemotype analogue in *L*_*j*_ was similar below a certain Tanimoto distance threshold was plotted against that threshold, to obtain a cumulative distance function (CDF). This was also performed for a reverse comparison of chemotypes in *L*_*j*_ compared to *L*_*i*_. Representative chemotypes for each cluster were represented dynamically as the minimally distant ligand pair between *L*_*i*_ and *L*_*j*_, for each cluster.

To emphasize highly similar ligands and de-emphasize lower similarity occurrences, CDFs were weighted according to the exponentially decaying weight function, 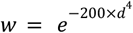, where ***w*** and ***d*** are weighted and unweighted Tanimoto distances, respectively, and ***e*** is the exponent, Euler’s number^23^ (**Supplementary Fig. 2**). Finally, the AUC of each ligand set pair CDF was normalized by the AUC of the reference CDF of the exponentially decaying weight function to calculate the similarity scores *S*_*ij*_ and *S*_*ji*_. These scores represent the number of unique chemotypes shared between *L*_*i*_ and *L*_*j*_.

#### Assignation of pharmacologically similar and dissimilar GPCR pair labels

The pharmacological similarity scores *S*_*ij*_ and *S*_*ji*_ were used to classify GPCR pairs as pharmacologically similar, dissimilar, or ambiguous (gray zone). We designed a hyperbolic generic formula with interpretable terms to facilitate fuzzy-like thresholds for *S*_*ij*_ and *S*_*ji*_ scores (details in **Supplementary Note 1**):

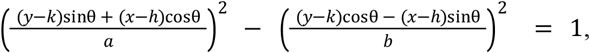

where ***x*** and ***y*** are the –log_10_-transformed *S*_*ij*_ and *S*_*ji*_ values to be substituted into the equation, ***θ*** is the angle of rotation in radians of each point (***x, y***) around the center (***h, k***), ***b*** defines the perpendicular distance between the vertex along the transverse axis to the asymptotes (and thus the curvature of the hyperbola), and ***a*** defines the distance between the vertex of the hyperbola and the centre, (***h, k***), of the branches of the hyperbola. Inequalities can then be used to define inclusion or exclusion criteria as appropriate.

In this study, we defined two hyperbolic inequalities, one defining dissimilar GPCR pairs with a maximum *S*_*ij*_ and *S*_*ji*_ score of 1 such that scores falling within the hyperbola were considered dissimilar, and another defining pharmacologically similar GPCR pairs with a maximum *S*_*ij*_ and *S*_*ji*_ score of 5 such that scores falling outside the hyperbola were considered similar. In simple terms, similar GPCRs shared approximately 5 chemotypes, whilst dissimilar GPCRs shared less than approximately 1 between them.

#### Application of pharmacological similarity/dissimilarity classifications to evaluate predictive performance via ROC curves

To compare the performance of TM or binding pocket similarity against any of the GPCR-CoINPocket variants, receiver operating characteristic (ROC) curves were generated. Pharmacologically similar pairs were defined as true, and dissimilar pairs as false. Pairs were then assigned a similarity score generated by the predictive models in question (e.g., TM similarity), and for each similarity score cut-off the fraction of true values with similarity scores exceeding the cut-off (true positives) was plotted against the fraction of false values in the same subset (false positives). For each of the tested methods, the resultant ROC curve was scaled for early prediction by square-root transformation of the x-axis, and the area under the curve (AUC) was calculated.

### GPCR-CoINPocket Framework

#### Calculation of ligand contact strength profiles

For each Pocketome entry, ligand contact strength fingerprints were calculated as previously described^14-16^. Contact strengths were approximated using the interatomic distance ***d*** between non-hydrogen atoms of the ligand and protein as a computationally efficient surrogate for interaction energies, such that strengths were 1 for ***d*** < ***d***_***min***_ = 3.23 Å, 0 for ***d*** > ***d***_***max***_ = 4.63 Å, and linearly decreasing from 1 to 0 as a function of ***d*** for ***d***_***min***_ < ***d*** < ***d***_***max***_. Contact interaction types were classified as polar, non-polar, or backbone interactions using BaSiLiCo ^32^, which categorizes these interactions based on the part of the amino acid side chain interacting with the given fragment.

To generate GPCR-CoINPocket scores, contact strengths were first calculated across 40 unique Class A GPCR structures as previously described^13^. Score aggregation used only side chain strengths, apart from glycine for which backbone interactions were treated as side chain. Several measures were taken to eliminate redundancy and prevent the most well-studied receptors from dominating the identified contact pattern. Specifically, side chain contact strengths were averaged across multiple occurrences of the same ligand in each Pocketome entry, and weighted by a value ranging from 0 to 1. This value represented the inherent flexibility of the binding residue as determined by the distribution of its observed root mean squared deviation between pairs of structures within the given Pocketome entry. The inherent flexibility coefficient was 1 for all side chains in single-structure entries. For Pocketome entries with more than one unique ligand (e.g. β_2_ adrenoceptor), fingerprints were clustered and only residues making interactions with strength > 0.5 in the top 80% of the list contributed to the consensus fingerprint, to emphasize only highly relevant critical contacts. Finally, contact strength values were multiplied by their relative frequency within the Pocketome entry to provide a final consensus fingerprint associated with the given Class A GPCR Pocketome entry.

To visualize ligand contact fingerprints, per-pocket contact strength vectors were projected onto a GPCRdb numbering system^49^ (analogous to the Ballesteros-Weinstein numbering scheme^50^) and represented with dots, where each dot is proportional to the square-root of the positional contact strength for that pocket (**Extended Data Fig. 4**).

#### Calculation of GPCR-CoINPocket scores across all Class A GPCRs

A pairwise GPCR-CoINPocket score was calculated for every pair of GPCRs in a multiple sequence alignment of 285 Class A GPCRs. This score was informed by 40 contact strength fingerprints (or “pockets”) derived from unique Class A GPCRs in the Pocketome v18.04. Briefly, for each of the 40 ligand contact fingerprints, the MSA of 285 GPCRs was restricted to only the relevant binding positions, and each position in the MSA was given an associated contact strength based on the relevant contact fingerprint. For each of the 40 unique pockets, the contact strength fingerprint of the given pocket was multiplied, element-wise, by the amino acid similarities at each position of the alignment.

For the original GPCR-CoINPocket, these similarities were derived from the Gonnet amino acid similarity matrix ***M*** such that the pairwise similarity ***S*** between amino acids ***i*** and ***j*** was defined as 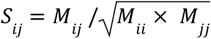. For GPCR-CoINPocket v2, similarities were optimized pairwise amino acid similarities from either the chemically-informed or unconstrained contact-informed amino acid similarity matrix. After weighting each position of the sequence similarities by the positional contact strengths, pairwise scores were summed, representing the similarity score for the GPCR pair from the perspective of the given pocket. This was performed for every unique Class A pocket, generating 40 sets of pairwise GPCR similarity scores. The differences in fingerprint sizes and pocket conservation distribution were handled by independent *z*-score transformations of each similarity set. The final GPCR-CoINPocket score for each GPCR pair is the average of the *z*-scored similarities across each of the 40 pocket-based similarities.

### Construction of Chemically-Informed Amino Acid Matrices

#### Ligand decomposition into commonly-represented ChEMBL-derived bioisosteres

A bioisostere fragment database of 83,043 fragments was provided by Molsoft LLC (La Jolla, CA, USA), and is freely available upon request. The database was derived from ChEMBL and informed by a global analysis of the ChEMBL activities across scaffolds with only one point of difference. In ICM, bioisostere fragments were substructure-matched to all small molecule ligands across 3,306 unique liganded protein ensembles in Pocketome v18.04^23^. Ligands were then replaced by their respective relevant fragments in ICM. Protein ligands were excluded from the analysis as the bioisostere fragment database did not include amino acid fragments and retention of the whole peptide would limit representation across structures.

#### Pairwise amino acid comparisons using bioisostere fragment:amino acid contact and non-contact vectors

Contact strengths for each fragment interacting with each residue across all structures in Pocketome were measured as described earlier and classified as polar, non-polar, or backbone interactions using BaSiLiCo. Contacts were recorded once per position, whilst non-contacts were recorded once per pocket so as to limit skew towards non-contacts. To ensure analysis proceeded only on appropriately-sized vectors, fragments were only considered if they made at least 10 contacts in total across any structure with any amino acid. The resultant 3,984 fragments were further condensed into meaningful vectors to control for redundancy by bottom-up hierarchical clustering using the UPGMA algorithm as implemented in ICM, with clusters defined at a Tanimoto distance cut-off of 0.1. This resulted in 1,233 bioisostere clusters, representing 1,233 vectors, for a final interaction dataset of contact (non-zero strength) and non-contact (zero strength) interaction vectors with the dimensions of 1,233 bioisostere clusters × 20 amino acids × 3 contact types (polar, non-polar, and backbone).

Pairwise amino acid differences were calculated for each contact type (polar, non-polar, backbone) using each of the 1,233 contact and non-contact vectors. A sequential scoring schema was used to assess: First, whether a fragment cluster interacted proportionally more with one amino acid over another using a chi-squared test, and if not (p-value > 0.05), then; Second, whether it interacted more strongly with one amino acid over another using a non-parametric Mann-Whitney/Wilcoxon rank-sum t-test. For chi-squared testing, contact and non-contact vector sizes were used to build a two-way contingency table for each cluster and contact type, and p-values simulated by Monte Carlo sampling of 2000 tables from a set of contingency tables with the same marginals, as implemented in R. For clusters deemed insignificant (p-value > 0.05), contact strength distributions in contact vectors were then compared using a Mann-Whitney/Wilcoxon non-parametric, rank-sum t test as implemented in R, and the subsequent p-value recorded irrespective of a significant result.

Comparisons of non-polar amino acids (Pro, Gly, Ala, Val, Ile, Leu, Met, or Phe) against any of the polar amino acids (Cys, Asp, Glu, Lys, Arg, or His) were manually set to the minimum p-value across the entire polar amino acid comparison dataset to denote strong differences. Non-polar amino acids compared against themselves in polar calculations had p-values set to the maximum p-value of the dataset to denote high self-to-self similarity. Where Gly was compared to any amino acid in non-polar comparisons, the value was set to the minimum p-value.

To calculate the overall difference score for each amino acid pair, the kernel-smoothed densities of –log_10_-transformed p-values of all 1,233 clusters were generated for each amino acid pair, for each contact type. The AUC of the significant fraction of each density distribution (values ≥ –log_10_(0.05)) was taken as a fraction of the total AUC of each density plot to obtain the final fractional difference score for each amino acid pair. Pairwise amino acid difference scores for each contact type were used to generate 3 amino acid comparison matrices informed by polar, non-polar, or backbone contact patterns.

For function optimization, these difference values were subtracted from 1 to generate similarity scores, as necessary for incorporation into GPCR-CoINPocket.

### Matrix Optimization

#### Optimization of a chemically-informed amino acid similarity matrix

The three component amino acid matrices (polar, non-polar, and backbone) were combined into a single chemically-informed amino acid similarity matrix. A function to combine component polar, non-polar, and backbone-informed amino acid similarity matrices to create a final chemically-informed matrix M was designed such that:

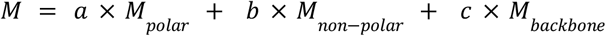

where ***a, b***, and ***c*** are coefficients to be optimized, representing the relative contribution of each matrix to the final matrix. Constraints were set such that ***a*** and ***b*** were positive, and the sum of ***a, b***, and ***c*** was 1.

The weight coefficients were optimized against the benchmark of pharmacologically similar/dissimilar GPCR pairs designed in this study, using the gradient-free Nelder-Mead simplex and Constrained Optimization BY Linear Approximation (COBYLA) algorithms as implemented in the *SciPy*^*51*^ package (v1.7.2) in Python 3.9. The cost function output to be minimized was 100 minus the AUC of an ROC curve with a square-root-transformed x-axis to emphasize the early recognition of chemically similar Class A GPCR pairs. To ensure that the final chemically-informed amino acid matrix would be educated by only the most important and relevant contacts, the ROC curves were generated by passing the combined matrices generated by the optimizer searches through GPCR-CoINPocket, replacing the Gonnet amino acid comparison matrix.

For both COBYLA and Nelder-Mead simplex methods, only polar and non-polar weight coefficients were exposed, with backbone weights derived as 1 minus the sum of polar and non-polar weight coefficients. Using COBYLA, constraints were set such that coefficients were ranged from 0 to 1 and summed to 1 to represent fractional similarities. Only the range constraint could be enforced with the Nelder-Mead simplex algorithm. A grid search of polar and non-polar weight coefficients was employed to systematically search all possible polar and non-polar weight coefficient combinations in (0, 1)^2^ in steps of 0.1, generating 121 start points each for both COBYLA and Nelder-Mead simplex methods. The algorithms were stopped upon convergence, defined by ≤ 0.001 change in the ROC curve AUC upon successive minimizations.

To assess the stability of the weight constraints (**Extended Data Fig. 5**), a stratified 5-fold cross-validation approach was used. The dataset of pharmacologically similar and dissimilar receptor pairs was partitioned into five equal subsets (folds), ensuring each fold maintained the same proportion of similar and dissimilar pairs. In each of the five iterations, four folds were used to train the weights, while the remaining fold served as the validation set. This process was repeated until every fold had been used exactly once for validation, providing a comprehensive assessment of weight stability.

#### Optimization of an unconstrained contact-informed amino acid similarity matrix

The optimization of the unconstrained contact-informed amino acid similarity matrix was conducted similarly to that of the chemically-informed amino acid similarity matrix, with a few differences. The Nelder-Mead simplex algorithm was used to optimize the amino acid similarities directly, without incorporation of chemical contact matrices (polar, non-polar, backbone). Further, each of the 190 possible pairwise amino acid combinations were allowed to vary independently between −1 and 1, except for self-to-self comparisons which were fixed at 1. The value range was chosen to conceptualize penalties for highly dissimilar amino acids (−1), indifference (0), and identity (1).

To imitate a grid-like approach, optimization was initialized from 380 different start points. In 190 start points, all weights were initialized randomly between 0.9 and 1.0, except a different amino acid pair was initialized between −1 and −0.9 each time. The opposite approach was used for the other 190 start points. As before, optimizations ceased when incremental improvements in the square-root-transformed x-axis ROC AUC were ≤ 0.001, approximating local minima, and amino acid pairs were educated by only the most important and relevant contacts through GPCR-CoINPocket.

### Experimental and Computational Validation of GPCR-CoINPocket

#### Experimental compounds and materials

Experimental chemicals and reagents were purchased from Sigma-Aldrich (Castle Hill, New South Wales, Australia), unless otherwise stated. Experimental compounds were purchased from Tocris Bioscience (Noble Park North, Victoria, Australia), unless otherwise stated. All compounds were >98% pure as determined by the manufacturer. General laboratory and tissue culture plasticware was purchased from Corning (Pennant Hills, New South Wales, Australia). Human β_2_ adrenoceptor in pcDNA3.1 was obtained from the cDNA Resource Centre (cat #AROB200000).

#### Cell culture

COS-1 cells were obtained from American Type Culture Collection (CRL-1650). Cells were maintained in Dulbecco’s Modified Eagle Media (DMEM) supplemented with 10% fetal bovine serum (FBS), incubated at 37°C and 5% CO_2_, and passaged using standard sterile tissue culture technique.

#### Cell membrane preparation

COS-1 cells were plated at a density of 4×10^6^ cells/flask into T175 flasks and incubated for 16 hours at 37°C and 5% CO_2_ in DMEM supplemented with 10% heat-inactivated FBS. After incubation, cells were transiently transfected with β_2_ adrenoceptor using PEImax 40K (Polysciences, Warrington, Pennsylvania, USA). Briefly, a transfection cocktail was made by mixing 15 µg/flask of β_2_ adrenoceptor-containing plasmid and 3 mL/flask of 150 mM NaCl, after which 60 µL/flask of PEImax was added to the cocktail and briefly vortexed. Following incubation a 20 min incubation at room temperature, 3 mL of the transfection cocktail was added to each flask, and supplemented with 10% FBS/DMEM up to 20 mL. Cells were incubated at 37°C and 5% CO_2_ for 24 hours, after which they were replated into 15 cm dishes and returned to the incubator. After 48 hours, cell membranes were harvested.

For the harvest of cell membranes, all steps were performed on wet ice using ice-cold reagents unless otherwise stated. Each 15 cm dish was gently rinsed with 10 mL of 1× phosphate-buffered saline (PBS) three times. Cells were then mechanically scraped into 5 mL/plate of 1x PBS and centrifuged at 500×g for 5 minutes. The supernatant was discarded, and the pellet resuspended in 20 mL of 20 mM HEPES/10 mM EDTA. The cell suspension was incubated on ice for 3 minutes to facilitate cell expansion in the hypotonic buffer before homogenising with an Ultra-TURRAX in 4×15-second bursts. The homogenized suspension was centrifuged at 600×g for 10 minutes, and the pellet discarded. The supernatant was centrifuged at 40,000×g for 1 hour to pellet cell membranes. After discarding supernatant, membrane pellets were resuspended in 50 mM Tris/2 mM EDTA/12.5 mM MgCl_2_, pH 7.4, and re-homogenized using a 31-gauge 1 mL insulin syringe before aliquoting and storing at –80°C for later use. The protein concentration was determined by comparison against a standard bovine serum albumin curve using Bradford reagent (Thermo Fisher Scientific, North Ryde, New South Wales, Australia).

#### Selection of experimental ligands

The top 10 possible pharmacological neighbor GPCR peptide families as ranked by their GPCR-CoINPocket v2 scores using the chemically-informed amino acid similarity matrix were considered for ligand picking. To select ligands for experimental validation, a database of relevant purchasable small molecules was obtained from Tocris Bioscience. Ligand sets from relevant receptors were clustered by Tanimoto distance in a bottom-up agglomerative hierarchical approach using the UPGMA algorithm as implemented in ICM, with cut-offs automatically set by ICM ranging from 0.2 for ligand sets with high internal similarity to 0.4 for ligand sets with high internal divergence. Chemical centres from these clusters were selected in ICM and a final 8 ligands were selected to maximize cost-efficiency and coverage over chemical diversity and receptor types.

#### Experimental ligand preparation

All experimental compounds were stored as solubilized aliquots at −20°C until required, except propranolol, which was prepared freshly in Milli-Q® water at a stock concentration of 10 mM for every experiment, and stored on wet ice until required. J 2156, TC-G 1003, SHA 68, and JNJ 10397049 were solubilized in DMSO and stored at 50 mM stock concentrations. WAY 267464 was solubilized in DMSO and stored at a stock concentration of 44 mM. ATC 0175 was solubilized in DMSO and stored at a stock concentration of 10 mM. GW 803430 was solubilized in 1 M hydrochloric acid and stored at a stock concentration of 20 mM. L-368,899 was solubilized in DMSO and stored at a stock concentration of 5 mM. TC-G 1003, WAY 267464, SHA 68, and JNJ 10397049 required heating at 45°C for 1 hour to solubilize at the highest working concentration but were soluble in binding buffer at final experimental concentrations.

#### Radioligand binding assays and affinity determination

Binding assays were performed using 1-5 µg crude COS-1 membranes transiently expressing β_2_ adrenoceptor in a total volume of 500 µL of 50 mM Tris/2 mM EDTA/12.5 mM MgCl_2_, pH 7.4 at room temperature. Membranes were defrosted on ice and re-homogenized with a 31-gauge 1 mL insulin syringe before use. Non-specific binding was defined using 10 µM of propranolol. Assays were terminated in a Brandel harvester by flushing four times with 4 mL of ice-cold 1x PBS and vacuum filtration through Whatman GF/C glass fibre filters. After drying, filters were incubated with 4 mL of Ultima GOLD scintillation cocktail for 1 hour in the dark at room temperature, before radioactivity was determined using a Perkin Elmer TriCarb-2800 TR scintillation counter. For saturation binding assays, membranes were incubated with increasing concentrations spanning 0.125-8 nM [^3^H]dihydroalprenolol ([^3^H]DHA) for 1 hour. For competitive binding assays, membranes were incubated with increasing concentrations of unlabelled competitor ligands and 0.5 nM of [^3^H]DHA for 1 hour, as indicated. For dissociation assays, membranes were equilibrated with 0.5 nM [^3^H]DHA for 1 hour at room temperature prior to initiating dissociation by the addition of 10 μM propranolol in the presence or absence of unlabelled inhibitors at various time points as indicated. Unlabelled competitor ligands were solubilized in water (propranolol), 1 M HCl (GW 803430), or DMSO (all other experimental compounds), with final concentrations of up to 2.2% DMSO or 5% 1 M HCl as appropriate. GraphPad Prism v9 was used to fit saturation, competition, and dissociation binding data to single site binding models and derive ligand affinities.

#### UMAP visualization of Class A GPCR clusters

For UMAP, pharmacological similarity and dissimilarity definitions were derived from ChEMBL v35. To visualize Class A GPCR clusters with UMAP, each GPCR was represented as a vector of its GPCR-CoINPocket scores against all other GPCRs, for each representative pocket. Representative unique pockets were generated by expanding each of the 40 pocket ligand contact fingerprints to the full size of all pockets to generate accurately aligned fingerprints, then by agglomerative bottom-up hierarchical clustering of the interaction vectors by UPGMA as implemented in ICM. Distance cutoffs were set to 0.35, producing 29 representative pockets. For each pocket, the pairwise GPCR similarities were generated in accordance with GPCR-CoINPocket v2 using the unconstrained contact-informed matrix. UMAP, as implemented in the R package *umap*, was then used to project GPCRs to two dimensions for visualization of the clusters.

To estimate the quality of cluster projection, all pairwise euclidean distances between each possible GPCR pair were measured and plotted as a distribution for pharmacologically similar or dissimilar pairs, as classified earlier in this study according to ChEMBL v27.

#### Prediction of receptor-ligand complex geometries

Binding sites and poses for the experimentally confirmed ligands in the β_2_ adrenoceptor were predicted using Boltz-1^16,38^ and AlphaFold 3^14^ (v3.0.1), both locally installed on the UCSD Triton Shared Computing Cluster. The amino-acid sequences and ligand SMILES strings used for modeling are provided as **Supplementary Data 5**. Models were generated using one random seed and 5 models per seed. All other parameters for both Boltz-1 and AlphaFold 3 were as default. Model inter-chain ipTM (interface predicted Template Modeling) score and minimum interchain PAE (predicted aligned error) were extracted from the summary json files returned by the predictors. Models were further visualized and analyzed in ICM v3.9-4a^52^ (Molsoft LLC) and both the full-atom^53^ and RTCNN^54,55^ scores were calculated. Analyzed co-folded models are provided as **Supplementary Data 6**.

## Data Availability

The molecular coordinates for all protein:ligand complexes used to generate ligand interaction fingerprints informing the amino acid contact-type matrices are part of Pocketome v18.04 on Zenodo, available at [link to be added]. Aggregated bioisostere cluster-to-residue interaction vectors and cluster member SMILES have been provided as **Supplementary Data 3**.

## Code Availability

BaSiLiCo is available as a public repository on GitHub: https://github.com/Kufalab-UCSD/BaSiLiCo. Code required for the measurement amino acid and ligand interaction frequency and strength from BaSiLiCo-labelled protein:ligand complexes, statistical scoring to generate polar-, nonpolar-, and backbone-based amino acid difference matrices, ChEMBL mining and generation of similarity benchmarks, and optimization of GPCR-CoINPocket are available as a public repository on GitHub: [to be added].

## Acknowledgements

We thank: Brendan P. Wilkins. and Erica Leonar for provision of COS-1 cell membranes; MolSoft LLC staff scientists for assistance with ICM queries and generous support with licensing extensions; and members of the Smith and Kufareva labs for productive and critical discussion of the work.

This work was supported by funding from the National Heart Foundation Future Leader Fellowship Level 2 (N.J.S.), the Simon Lee Foundation Scholarship (S.S.S.), the Australian Government Research Training Program (S.S.S. and T.N.), the National Health and Medical Research Council C.J. Martin Early Career Fellowship 1145746 (T.N.), the National Institute of Health R01 Grants GM136202, AI161880, and AI118985 (I.K.), the National Institute of Health R21 Grants AI149369 and AI156662 (I.K.), the University of California Office of the President (UCOP) Cancer Research Coordinating Committee (CRCC) seed grant C26CR10041 (I.K.), and the CSD/UCSF Cancer Cell Mapping Initiative Pilot Grant under NIH U54 CA274502 (I.K.).

## Author Contributions

S.S.S., N.J.S., T.N., and I.K. conceived, designed, and supervised the study. S.S.S. and I.K. performed computational experiments. A.M.F. supervised S.S.S. in radioligand binding and provided membranes for experiments. S.S.S., N.J.S., and I.K. analyzed results. R.P.R. manually curated Class A GPCR alignments. R.A. supported S.S.S. with complex chemoinformatics. S.S.S., T.N., A.V.I., and I.K. wrote code for various components of the study. S.S.S., N.J.S., and I.K. wrote the manuscript. All authors contributed to editing the manuscript.

## Extended Data Figures

**Extended Data Fig. 1.**
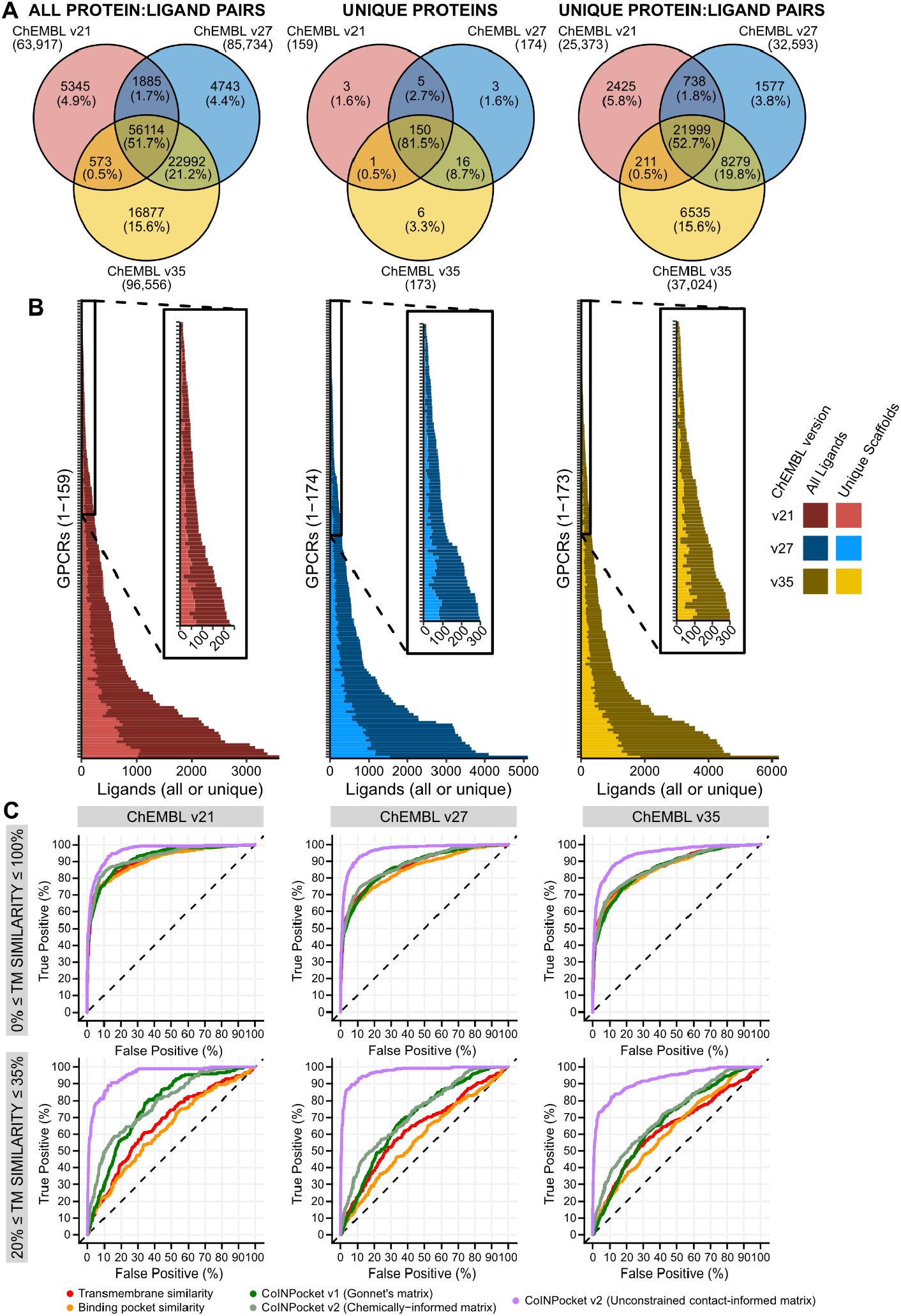
Comparison of ChEMBL versions 21, 27, and 35. **a**, Venn diagrams showing overlap of ChEMBL versions 21, 27, and 35 for various parameters, calculated using Class A GPCR ligand sets in a pairwise manner. **b**, Number of ligands or unique chemotypes for each Class A GPCR in ChEMBL versions 21, 27, or 35. Insets, zooms of GPCRs with few ligands. **c**, Comparison of GPCR-CoINPocket v1, transmembrane (TM) similarity, and binding pocket similarity in distinguishing pharmacologically similar GPCRs from dissimilar ones across either all GPCRs or those with homology too low for accurate modeling but sufficient similarity for high accuracy sequence alignment. Similar/dissimilar pairs were derived from ChEMBL versions 21, 27, or 35 as indicated.

**Extended Data Fig. 2.**
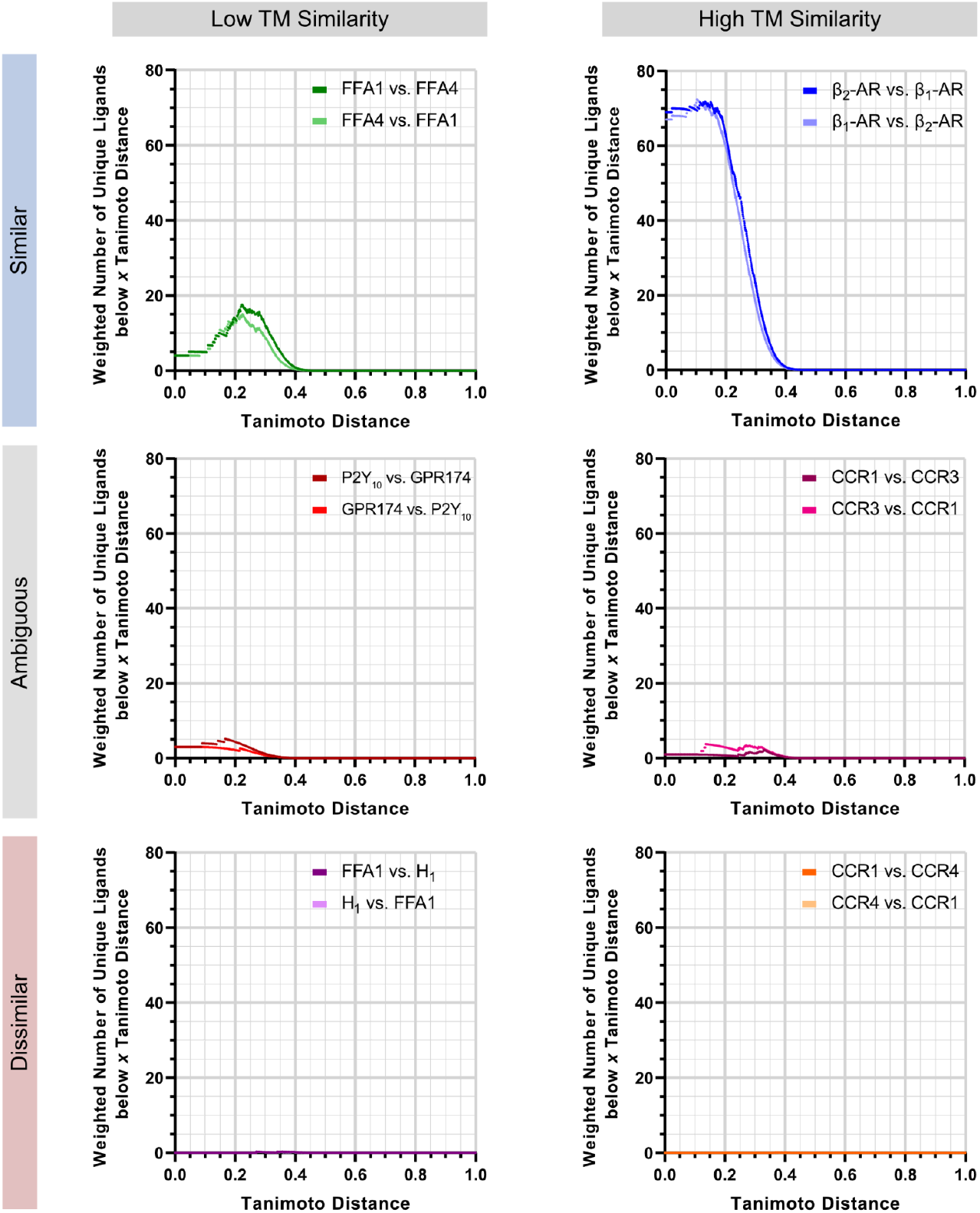
The hyperbolic classification function correctly captures known pharmacological similarities and dissimilarities of reference receptor pairs. The cumulative distribution function of the Tanimoto distances of unique chemotypes for each receptor pair were weighted using the exponential decay function as shown in **Supplementary Fig. 2**. The hyperbolic classification function correctly captured known similarities, ambiguities, and dissimilarities between weakly or highly homologous receptors. Data was sourced from ChEMBL v27. Receptors are named according to IUPHAR/Guide to Pharmacology standards.

**Extended Data Fig. 3.**
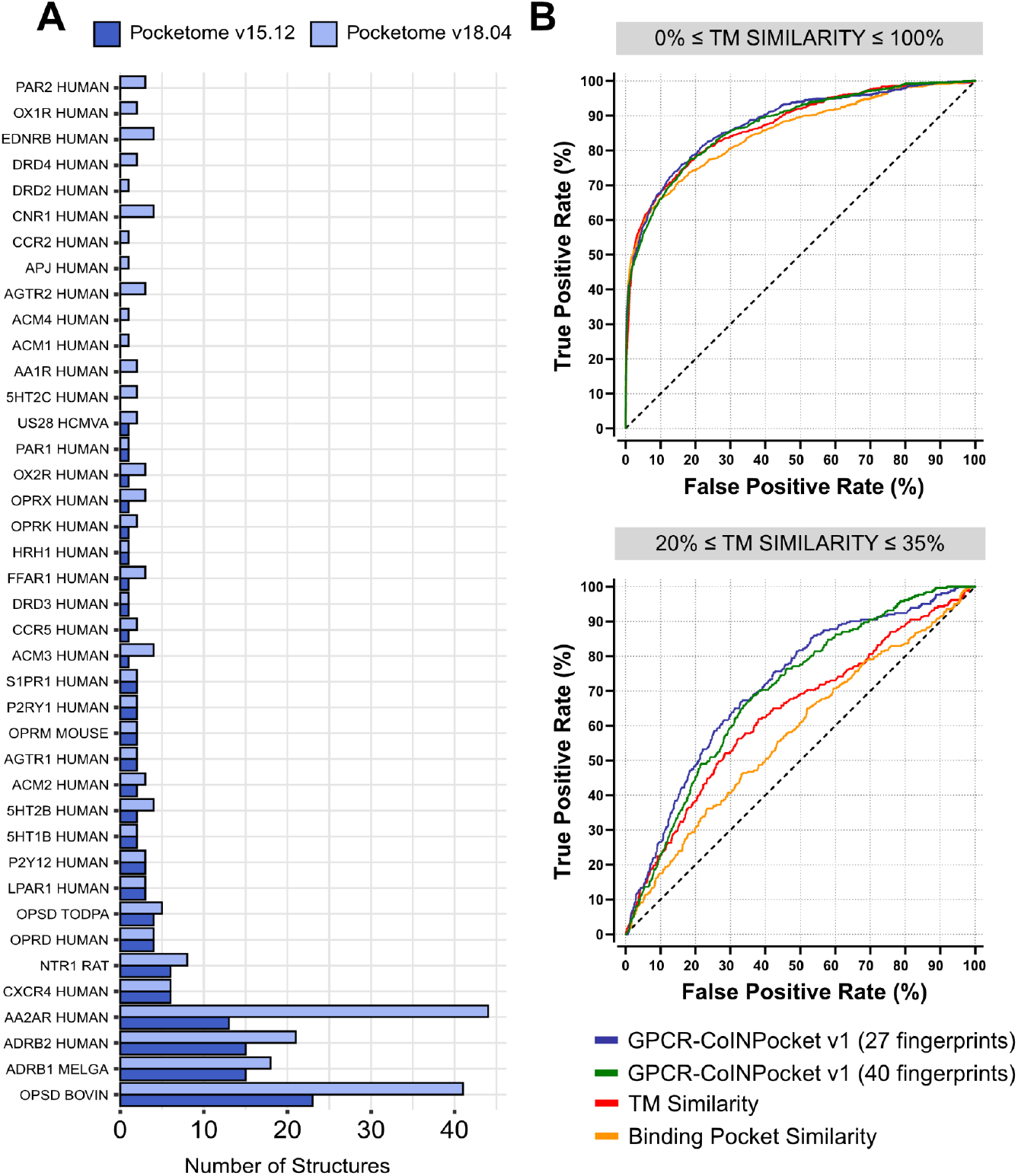
The impact of updated fingerprints on the predictive power of GPCR-CoINPocket and other predictive metrics. **a**, The distribution of Class A GPCRs in Pocketome v18.04 was compared to that of Pocketome v15.12, which was used for the original GPCR-CoINPocket implementation^13,23^. Receptors are named in accordance with UniProt nomenclature. **b**, ROC curves comparing GPCR-CoINPocket v1 using the original ligand fingerprints derived from 27 liganded Class A structures^13^, an updated set including 40 fingerprints, transmembrane (TM) similarity, and binding pocket similarity in distinguishing similar GPCR pairs from dissimilar ones across either all GPCRs or those with homology too low for accurate modeling but sufficient similarity for high accuracy sequence alignment. The benchmarking set used was derived from ChEMBL v27, as described in **Fig. 1**.

**Extended Data Fig. 4.**
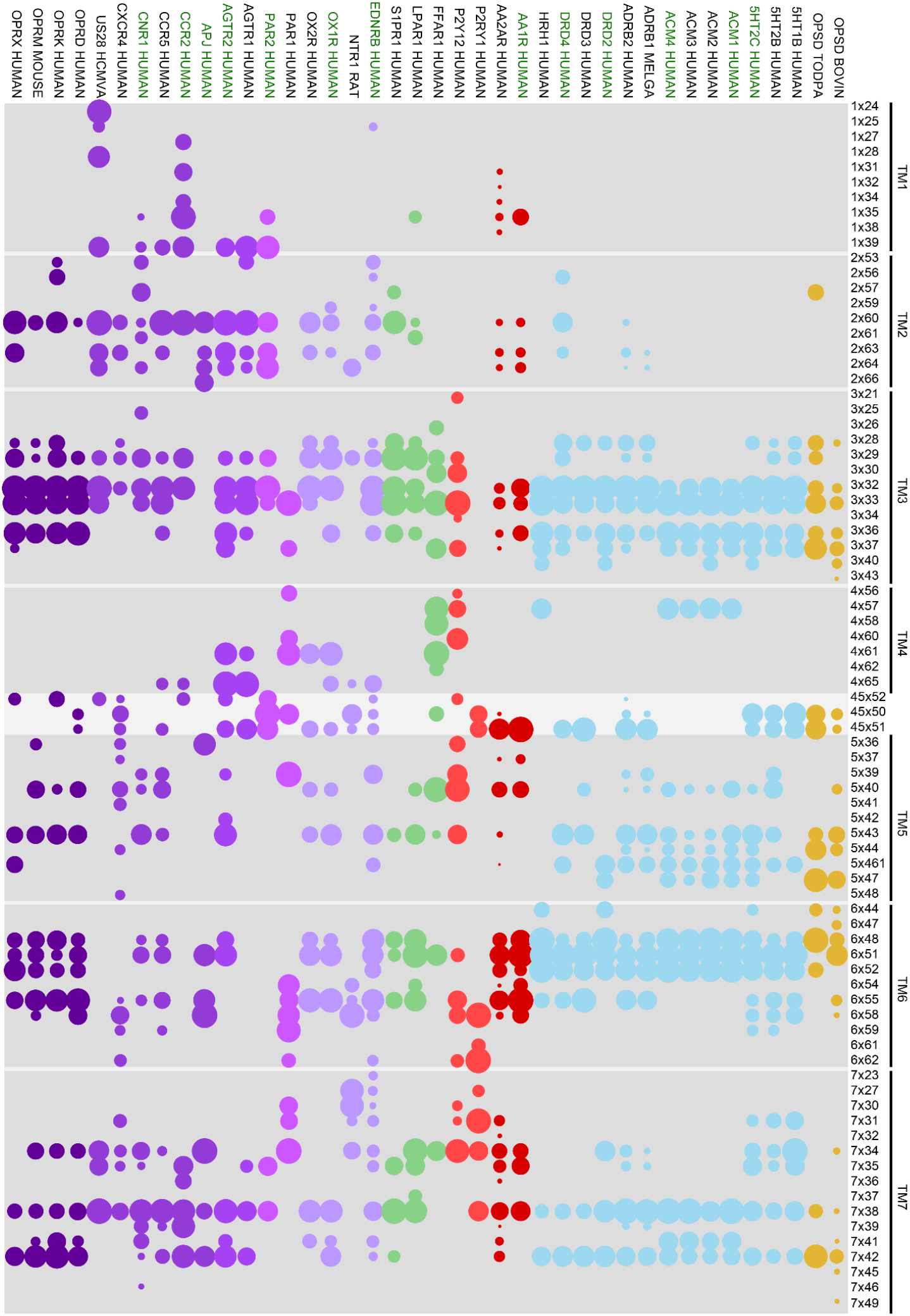
Class A GPCR ligand contact fingerprints. Ligand contact strengths were measured from 40 distinct GPCRs annotated in the latest Pocketome (v18.04)^23^. Circle sizes represent relative side chain contact strength and residue positions are numbered as per the GPCRdb numbering scheme^49^. Circle colors represent the chemical class of the receptor’s endogenous ligand: purple, peptide; green, lipids, red, nucleosides; blue, aminergics; gold, retinal. Contacts from loop regions were excluded, other than those within one residue of the conserved cysteine in ECL2. Weak “pinprick-sized” contacts were also removed to emphasize only highly relevant contact fingerprint patterns for this visualisation. Receptors are named according to UniProt nomenclature, and colored green for new entries compared to Pocketome v15.12 which was used for informing GPCR-CoINPocket v1.

**Extended Data Fig. 5.**
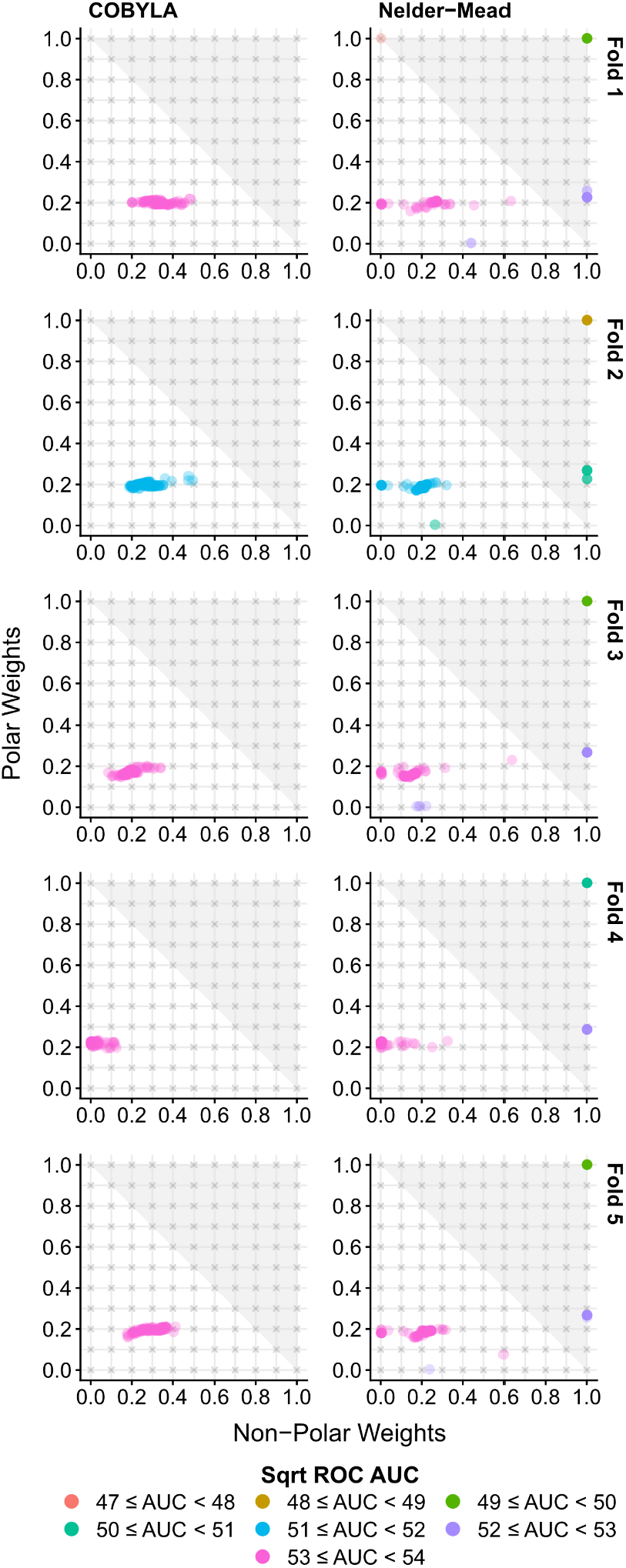
Consensus weights optimized by the Nelder-Mead simplex and COBYLA optimizers are stable. A scatterplot of the final minimized weight coefficients (non-polar and polar) exposed to the optimization algorithms. For each start point, a stratified 20% True/False dropout was applied. Gray crosses represent all start points used for both COBYLA and Nelder-Mead optimizers. The gray triangle represents disallowed regions where backbone weights are negative. Points represent final optimized weight coefficients. Point colors represent the early-detection-weighted ROC AUC of the final optimized weight triplet, including the derived backbone weight (1 – sum of non-polar and polar weights).

**Extended Data Fig. 6.**
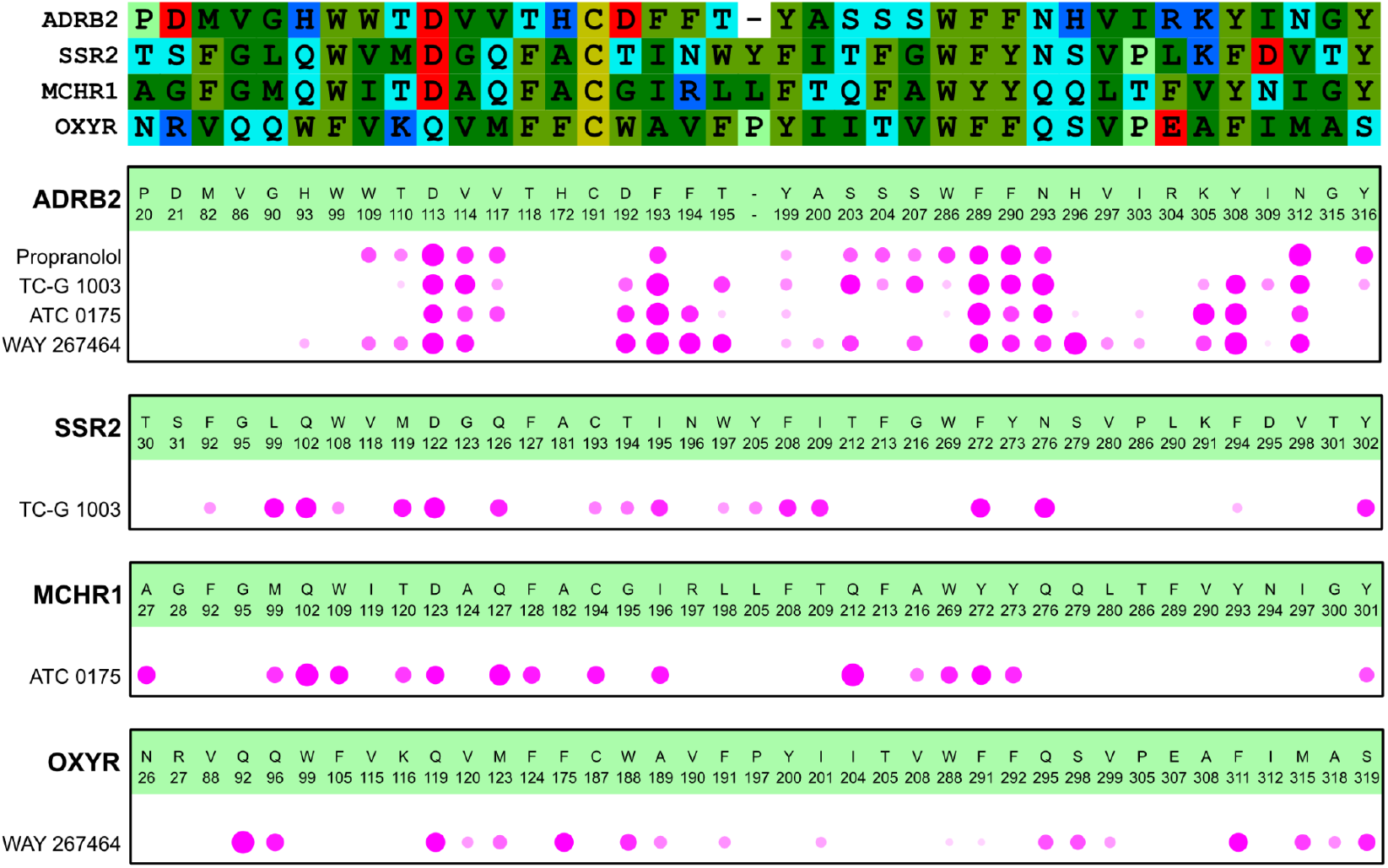
Contact fingerprints of hit compounds. **a**, The sequence alignment for the binding pocket defined by the interactions of propranolol and the hit compounds TC-G 1003, ATC 0175, and WAY 267464 against the β_2_ adrenoceptor and their cognate receptors is shown colored by the residue property: hydrophobic, dark green; aromatic, light green; proline, lime green; basic, blue; acidic, red; polar, cyan. **b**, The ligand contact fingerprints for these compounds against the β_2_ adrenoceptor and their cognate receptors is shown. Residue identities and numbers are displayed on top of each receptor panel. Pink circles indicate ligand contacts with the receptor, with size scaled to the strength of the contact. Fingerprints were measured using ICM v3.9-4a as described in **Methods** from top-scoring co-folded models.

**Extended Data Fig. 7.**
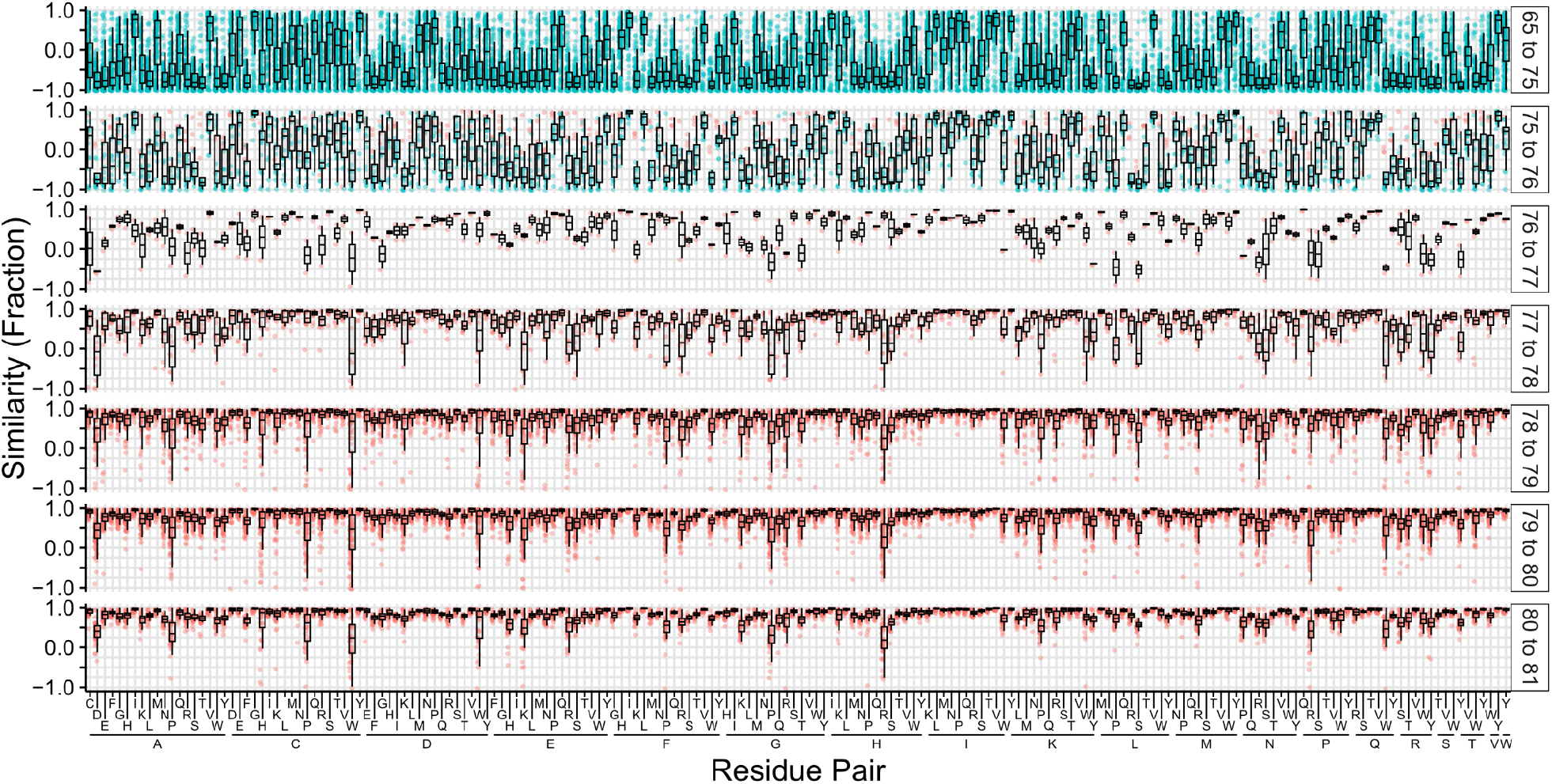
Optimization results for the unconstrained contact-informed amino acid similarity matrix. The individual optimized amino acid similarities are represented as a fraction for each pair, and the results for each optimization run are shown as individual points, colored by whether initialization was mostly positive (red) or negative (blue). Optimization results are faceted into square-rooted ROC AUC ranges for visual clarity. Boxplots are shown for each amino acid pair, indicating the flexibility of the optimization result for the given ROC AUC.

## Supplementary Information

**Supplementary Note 1: Derivation and application of the general form of a rotated hyperbola**

The generic formula of a hyperbola with point co-ordinates (***x, y***) around a central point (***h, k***) is as follows:

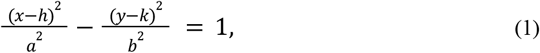

where ***b*** defines the perpendicular distance between the vertex along the transverse axis to the asymptotes (and thus the curvature of the hyperbola), and ***a*** defines the distance between the vertex of the hyperbola and the centre, (***h, k***), of the branches of the hyperbola.

To rotate points (***x, y***) around the centre (***h, k***), obtaining points (***x’, y’***), we use the following rotation matrix:

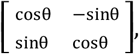

where ***θ*** is the angle of rotation around (***h, k***) in radians, such that:

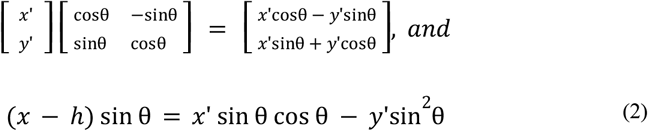

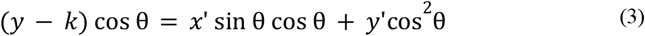

Subtracting (2) from (3):

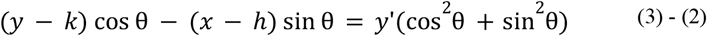

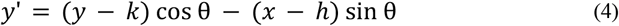

From (3):

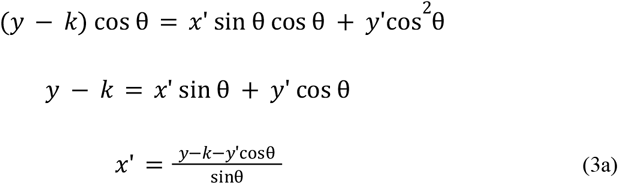

Substituting (4) into (3a) to solve for ***x’***:

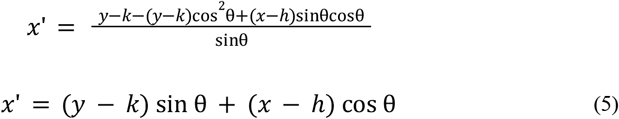

Combining (4) and (5), we arrive at the final generic equation for a hyperbola, rotated around the centre (***h, k***):

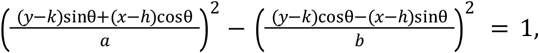

In this study, we described two hyperbolae to define pharmacologically similar or dissimilar GPCR pairs. For each of these hyperbolae, (***h, k***), the center, was dynamically defined as the maximum across all *S*_*ij*_ and *S*_*ji*_ scores. Parameter ***b***, the perpendicular distance between the vertex along the transverse axis to the asymptotes was defined as 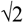 as a heuristic to accept as dissimilar cases when one of *S*_*ij*_ and *S*_*ji*_ scores were only slightly above 1. Finally, ***θ***, the angle of rotation around (***h, k***) in radians was defined as 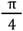(45°), since *S*_*ij*_ and *S*_*ji*_ scores were roughly correlated along ***y* = *x***.

For parameter ***a***, the distance between the vertex and center, we chose values corresponding to vertices for each of the hyperbolae to match an intuitive understanding of shared pharmacology. We reasoned that receptors with 1 or less shared scaffolds (*S*_*ij*_ and *S*_*ji*_ scores ≤ 1) should be considered dissimilar, corresponding to the origin (0, 0) on –log_10_ scale (**Fig. 1B**). Thus, ***a*** was defined as 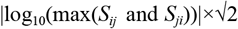. Conversely, we proposed that receptors with 5 or more shared scaffolds should be defined as pharmacologically similar, corresponding to a vertex at (–log_10_(5), –log_10_(5)). Thus, ***a*** was defined as 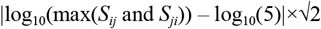.

Pharmacologically dissimilar or similar GPCRs could then be defined as inequalities < (within the boundary) or > (outside the boundary) of the two hyperbolae, respectively. We also note that the mirrored branch of each hyperbola does not play a role in classification, as the center of the branches is set such that the mirrored branch plots onto empty space.

## Supplementary Figures

**Supplementary Fig. 1.**
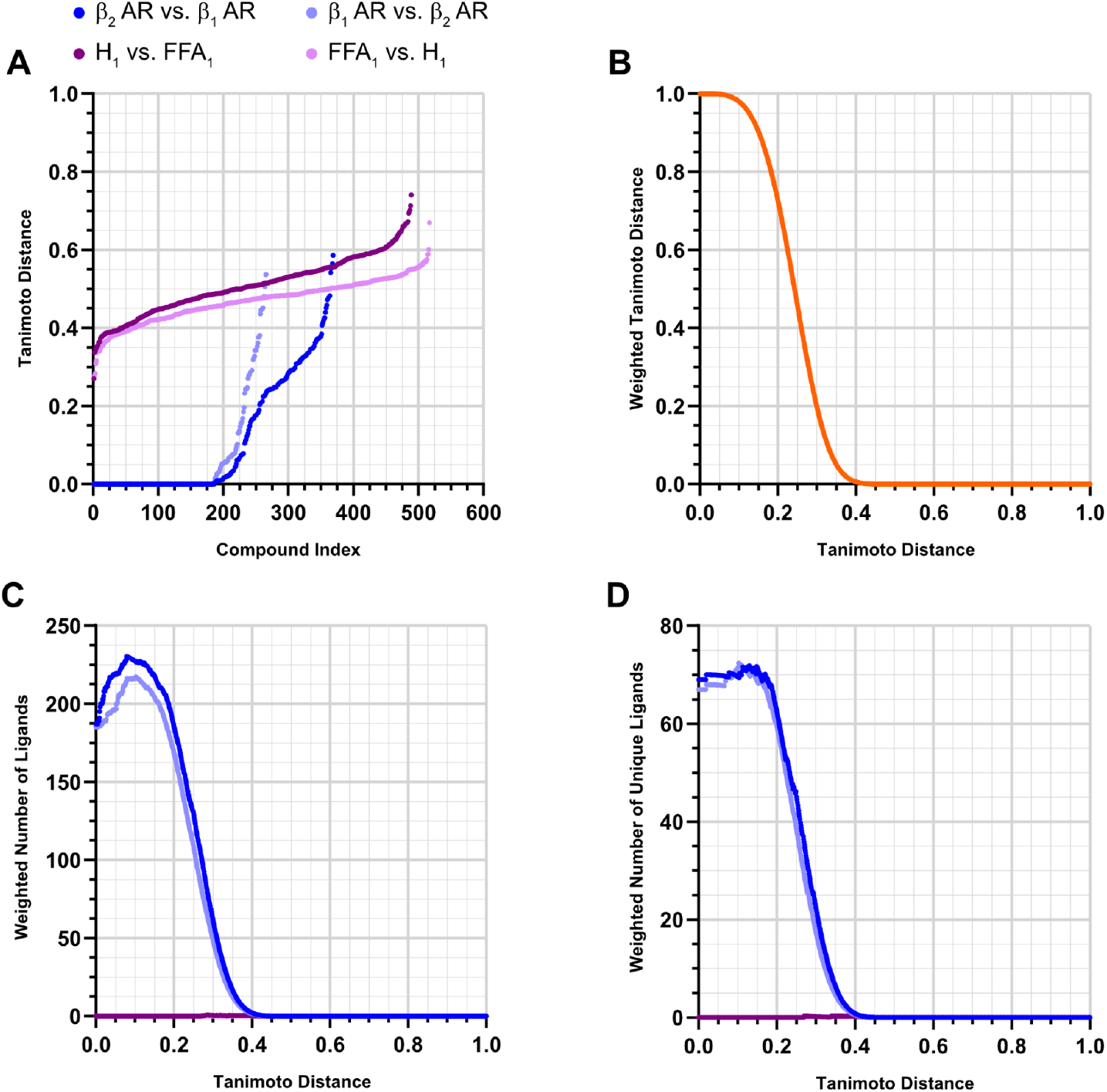
Refinements to the measurement of chemical similarity across ligand sets. A representative example of the application of the exponentially decaying weight function towards ligand sets is shown for a pharmacologically similar (β_1_ and β_2_ adrenoceptors) and pharmacologically dissimilar (FFA1 and H_1_ receptors) receptor pair, as defined by data from ChEMBL v27. **a**, The asymmetrical pairwise minimum unweighted Tanimoto distance for each compound is shown, ordered by increasing Tanimoto distance. **b**, The reference exponentially decaying weight function is plotted. **c**,**d**, Tanimoto distances from **a** were binned by cumulative distribution and shown (**c**) without or (**d**) with application of the weight function.

**Supplementary Fig. 2.**
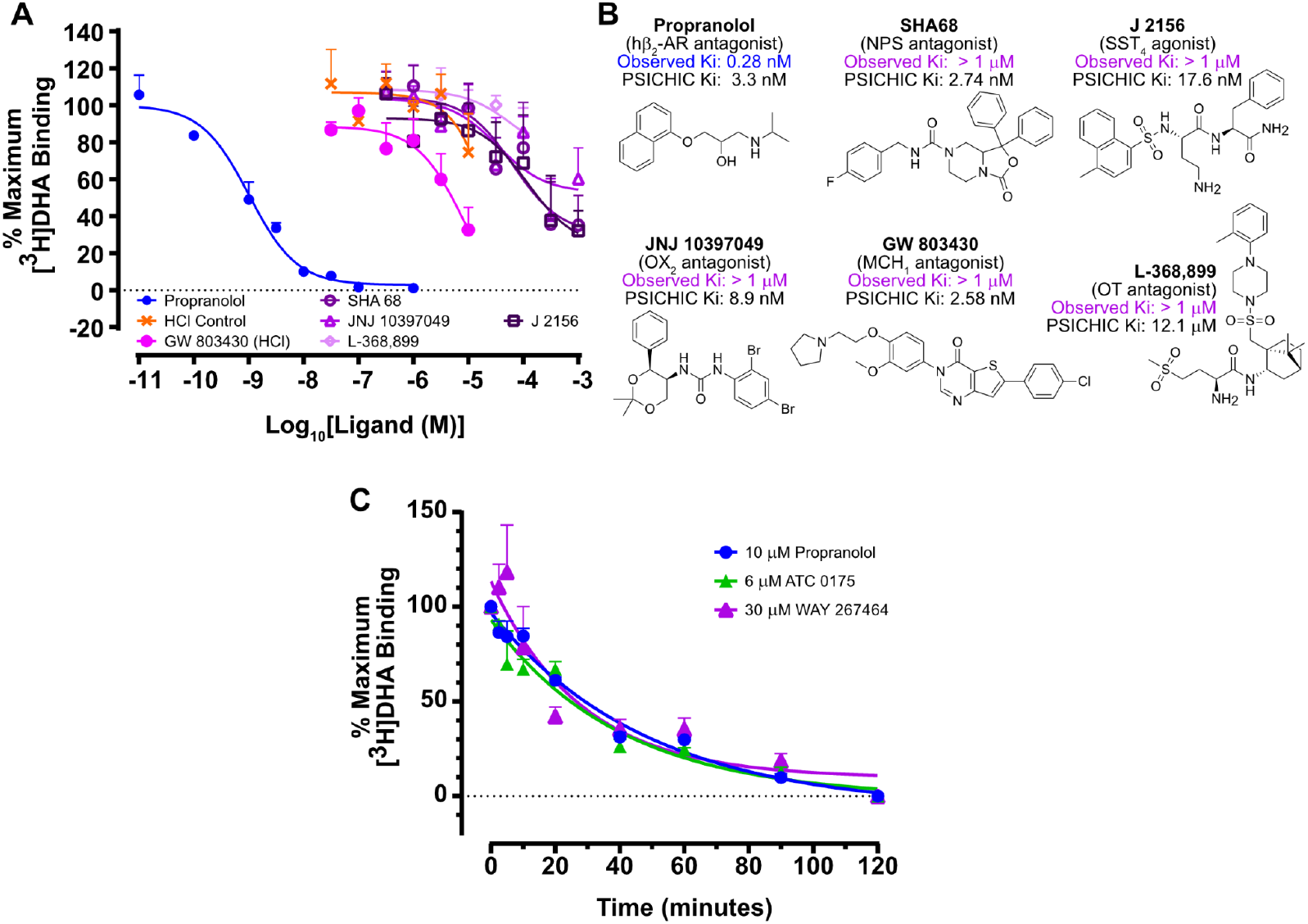
Competitive radioligand binding data for inactive compounds against the β_2_ adrenoceptor. **a**, Competitive radioligand binding assay using putative surrogate ligands. Binding assays used crude purified membranes of COS-1 cells transiently transfected with β_2_ adrenoceptor. Unlabelled competitor ligands were titrated against 0.5 nM [^3^H]DHA and their affinity derived. Points and bars represent means ± SEMs of n=3 experiments performed in triplicate. Curves were fit to a logistic single site binding model using GraphPad Prism v9. **b**, Chemical structures of weakly-binding or inactive compounds assessed in this study. Observed K_i_ (affinity) of experimental ligands from competitive radioligand binding > 1 μM (purple) or propranolol (blue) are displayed above each structure along with their PSICHIC-predicted^39^ K_i_. **c**, Radioligand dissociation assay of hit compounds in comparison to *bona fide* orthosteric antagonist propranolol. The rate at which [^3^H]DHA dissociated from the β_2_ adrenoceptor was measured over the course of 2 hrs at room temperature upon addition of 10 μM propranolol (blue) in the presence or absence of off-target compounds: 6 μM ATC 0175 (green) or 30 μM WAY 267464 (purple). Points and bars represent means ± SEMs of n=2-4 experiments performed in triplicate. Curves were fit to a logistic single site binding model using GraphPad Prism v9.

## Supplementary Data

**Supplementary Data 1:**

- Excel Spreadsheet: Pairwise amino acid similarities for each contact type (statistical scoring), for the chemically-informed matrix, and for the unconstrained contact-informed matrix.

**Supplementary Data 2:**

- Excel Spreadsheet: True/False classifications and pairwise receptor similarities (Pocket similarity, Transmembrane similarity, and CoINPocket scores: CoINPocket v1, CoINPocket v1 with updated fingerprints, CoINPocket v2 with the chemically-informed matrix, CoINPocket v2 with the unconstrained contact-informed matrix) using pharmacological similarities derived from ChEMBL v21, v27, or v35.

**Supplementary Data 3:**

- Zip file: Tables of bioisostere interaction and non-interaction vector strengths (both frequency-weighted and unweighted). Each file is a comma-separated file, with all backbone/polar/nonpolar interaction vectors recorded. From there, users may extract the frequencies easily. A key table defines which fragments are present in which clusters.

**Supplementary Data 4:**

- Sheet 1: β_2_ adrenoceptor neighbors scored by transmembrane similarity and CoINPocket v2 score (using the chemically-informed matrix and ChEMBL v27 definitions)
- Sheet 2: PSICHIC evaluation of β_2_ adrenoceptor with selected peptide receptor ligands, compared to radioligand binding data

**Supplementary Data 5:**

- Zip file: Amino acid sequences of receptors and compound SMILES for modeling

**Supplementary Data 6:**

- Zip file: ICM binary file (ICB) containing selected models presented in this study

## References

1. Hauser, A. S., Attwood, M. M., Rask-Andersen, M., Schiöth, H. B. & Gloriam, D. E. Trends in GPCR drug discovery: new agents, targets and indications. Nat. Rev. Drug Discov. 16, 829–842 (2017).

2. Alexander, S. P. H. et al. The Concise Guide to PHARMACOLOGY 2025/26: G protein-coupled receptors. Br. J. Pharmacol. 182 Suppl 1, S24–S151 (2025).

3. Laschet, C., Dupuis, N. & Hanson, J. The G protein-coupled receptors deorphanization landscape. Biochem. Pharmacol. 153, 62–74 (2018).

4. So, S. S., Ngo, T., Keov, P., Smith, N. J. & Kufareva, I. Tackling the complexities of orphan GPCR ligand discovery with rationally assisted approaches. in GPCRs (eds. Jastrzebska, B. & Park, P. S.-H.) 295–334 (Elsevier, 2020).

5. Huang, X.-P. et al. Allosteric ligands for the pharmacologically dark receptors GPR68 and GPR65. Nature 527, 477–483 (2015).

6. Lansu, K. et al. In silico design of novel probes for the atypical opioid receptor MRGPRX2. Nat. Chem. Biol. 13, 529–536 (2017).

7. Liu, C. et al. GPR139, an orphan receptor highly enriched in the habenula and septum, is activated by the essential amino acids L-tryptophan and L-phenylalanine. Mol. Pharmacol. 88, 911–925 (2015).

8. Szpakowska, M. et al. Inclusion of ACKR5 in the systematic nomenclature of atypical chemokine receptors. Nat. Rev. Immunol. 25, 225–226 (2025).

9. Chevigné, A. et al. International union of basic and clinical pharmacology. CXVIII. Update on the nomenclature for atypical chemokine receptors including ACKR5. Pharmacol. Rev. 77, 100012 (2024).

10. Meyrath, M. et al. ACKR5/GPR182 is a scavenger receptor for the atypical chemokine CXCL17, GPR15L and various endogenous peptides. bioRxiv 2024.06.01.596940 (2024) doi:10.1101/2024.06.01.596940.

11. Ocón, B. et al. A lymphocyte chemoaffinity axis for lung, non-intestinal mucosae and CNS. Nature 635, 736–745 (2024).

12. Ngo, T. et al. Orphan receptor GPR37L1 remains unliganded. Nat. Chem. Biol. 17, 383–386 (2021).

13. Ngo, T. et al. Orphan receptor ligand discovery by pickpocketing pharmacological neighbors. Nat. Chem. Biol. 13, 235–242 (2017).

14. Abramson, J. et al. Accurate structure prediction of biomolecular interactions with AlphaFold 3. Nature 630, 493–500 (2024).

15. Jumper, J. et al. Highly accurate protein structure prediction with AlphaFold. Nature 596, 583–589 (2021).

16. Wohlwend, J. et al. Boltz-1 democratizing biomolecular interaction modeling. bioRxivorg (2025) doi:10.1101/2024.11.19.624167.

17. Passaro, S. et al. Boltz-2: Towards accurate and efficient binding affinity prediction. bioRxivorg (2025) doi:10.1101/2025.06.14.659707.

18. Barril, X. Computer-aided drug design: time to play with novel chemical matter. Expert Opin. Drug Discov. 12, 977–980 (2017).

19. Gonnet, G. H., Cohen, M. A. & Benner, S. A. Exhaustive matching of the entire protein sequence database. Science 256, 1443–1445 (1992).

20. Dayhoff, M., Schwartz, R. & Orcutt, B. 22. A model of evolutionary change in proteins. in Atlas of Protein Sequence and Structure vol. 5 345–352 (1978).

21. Henikoff, S. & Henikoff, J. G. Amino acid substitution matrices from protein blocks. Proc. Natl. Acad. Sci. U. S. A. 89, 10915–10919 (1992).

22. Abagyan, R. A. & Batalov, S. Do aligned sequences share the same fold? J. Mol. Biol. 273, 355–368 (1997).

23. Kufareva, I., Ilatovskiy, A. V. & Abagyan, R. Pocketome: an encyclopedia of small-molecule binding sites in 4D. Nucleic Acids Res. 40, D535–40 (2012).

24. Becht, E. et al. Dimensionality reduction for visualizing single-cell data using UMAP. Nat. Biotechnol. 37, 38–44 (2018).

25. McInnes, L., Healy, J. & Melville, J. UMAP: Uniform manifold approximation and projection for dimension reduction. arXiv preprint arXiv:1802.03426 (2018).

26. Milligan, G., Alvarez-Curto, E., Hudson, B. D., Prihandoko, R. & Tobin, A. B. FFA4/GPR120: Pharmacology and therapeutic opportunities. Trends Pharmacol. Sci. 38, 809–821 (2017).

27. Yoshie, O. & Matsushima, K. CCR4 and its ligands: from bench to bedside. Int. Immunol. 27, 11–20 (2015).

28. Nomiyama, H., Osada, N. & Yoshie, O. A family tree of vertebrate chemokine receptors for a unified nomenclature. Dev. Comp. Immunol. 35, 705–715 (2011).

29. Inoue, A. et al. TGFα shedding assay: an accurate and versatile method for detecting GPCR activation. Nat. Methods 9, 1021–1029 (2012).

30. Kufareva, I., Katritch, V., Participants of GPCR Dock 2013, Stevens, R. C. & Abagyan, R. Advances in GPCR modeling evaluated by the GPCR Dock 2013 assessment: meeting new challenges. Structure 22, 1120–1139 (2014).

31. Abagyan, R., Chen, W. & Kufareva, I. Computational Approaches to Nuclear Receptors. 84–109 (Royal Society of Chemistry, Cambridge, England, 2012).

32. Ilatovskiy, A. V., So, S. S. & Kufareva, I. BaSiLiCo: Binding Site Ligand Contacts Visualization Tool. (Github: https://github.com/Kufalab-UCSD/BaSiLiCo).

33. Kufareva, I. et al. Status of GPCR modeling and docking as reflected by community-wide GPCR Dock 2010 assessment. Structure 19, 1108–1126 (2011).

34. Lin, H., Sassano, M. F., Roth, B. L. & Shoichet, B. K. A pharmacological organization of G protein-coupled receptors. Nat. Methods 10, 140–146 (2013).

35. Fredriksson, R., Lagerström, M. C., Lundin, L.-G. & Schiöth, H. B. The G-protein-coupled receptors in the human genome form five main families. Phylogenetic analysis, paralogon groups, and fingerprints. Mol. Pharmacol. 63, 1256–1272 (2003).

36. Fredriksson, R. & Schiöth, H. B. The repertoire of G-protein-coupled receptors in fully sequenced genomes. Mol. Pharmacol. 67, 1414–1425 (2005).

37. Gloriam, D. E., Foord, S. M., Blaney, F. E. & Garland, S. L. Definition of the G protein-coupled receptor transmembrane bundle binding pocket and calculation of receptor similarities for drug design. J. Med. Chem. 52, 4429–4442 (2009).

38. Mirdita, M. et al. ColabFold: making protein folding accessible to all. Nat. Methods 19, 679–682 (2022).

39. Koh, H. Y., Nguyen, A. T. N., Pan, S., May, L. T. & Webb, G. I. Physicochemical graph neural network for learning protein–ligand interaction fingerprints from sequence data. Nat. Mach. Intell. 6, 673–687 (2024).

40. Rabal, O., Castellar, A. & Oyarzabal, J. Novel pharmacological maps of protein lysine methyltransferases: key for target deorphanization. J. Cheminform. 10, 32 (2018).

41. Farhangdoost, N. et al. Chromatin dysregulation associated with NSD1 mutation in head and neck squamous cell carcinoma. Cell Rep. 34, 108769 (2021).

42. Topchu, I., Bychkov, I., Gursel, D., Makhov, P. & Boumber, Y. NSD1 supports cell growth and regulates autophagy in HPV-negative head and neck squamous cell carcinoma. Cell Death Discov. 10, 75 (2024).

43. Keiser, M. J. et al. Relating protein pharmacology by ligand chemistry. Nat. Biotechnol. 25, 197–206 (2007).

44. Hert, J., Keiser, M. J., Irwin, J. J., Oprea, T. I. & Shoichet, B. K. Quantifying the relationships among drug classes. J. Chem. Inf. Model. 48, 755–765 (2008).

45. Akinboye, E. S. & Bakare, O. Biological activities of emetine. Open Nat. Prod. J. 4, 8–15 (2011).

46. Michino, M., Vendome, J. & Kufareva, I. AI meets physics in computational structure-based drug discovery for GPCRs. 2, 16 (2025).

47. Chitsazi, R. et al. The 4thGPCR Dock: assessment of blind predictions for GPCR-ligand complexes in the era of AlphaFold. bioRxiv 2025.04.18.647407 (2025) doi:10.1101/2025.04.18.647407.

48. Yang, Y. et al. Efficient exploration of chemical space with docking and deep learning. J. Chem. Theory Comput. 17, 7106–7119 (2021).

49. Isberg, V. et al. Generic GPCR residue numbers - aligning topology maps while minding the gaps. Trends Pharmacol. Sci. 36, 22–31 (2015).

50. Ballesteros, J. A. & Weinstein, H. [19] Integrated methods for the construction of three-dimensional models and computational probing of structure-function relations in G protein-coupled receptors. in Methods in Neurosciences (ed. Sealfon, S. C.) vol. 25 366–428 (Elsevier, 1995).

51. Virtanen, P. et al. SciPy 1.0: fundamental algorithms for scientific computing in Python. Nat. Methods 17, 261–272 (2020).

52. Abagyan, R. & Totrov, M. Biased probability Monte Carlo conformational searches and electrostatic calculations for peptides and proteins. J. Mol. Biol. 235, 983–1002 (1994).

53. Neves, M. A. C., Totrov, M. & Abagyan, R. Docking and scoring with ICM: the benchmarking results and strategies for improvement. J. Comput. Aided Mol. Des. 26, 675–686 (2012).

54. Dawson, J. R. D. et al. Molecular determinants of antagonist interactions with chemokine receptors CCR2 and CCR5. bioRxivorg (2024) doi:10.1101/2023.11.15.567150.

55. Raush, E. et al. Novel GPU engines for virtual screening of Giga-sized libraries identify inhibitors of challenging targets. J. Chem. Inf. Model. 65, 10253–10268 (2025).

